# DNA double-strand break-capturing nuclear envelope tubules drive DNA repair

**DOI:** 10.1101/2023.05.07.539750

**Authors:** Mitra Shokrollahi, Mia Stanic, Anisha Hundal, Janet N. Y. Chan, Defne Urman, Anne Hakem, Roderic E. Garcia, Jun Hao, Philipp G. Maass, Brendan C. Dickson, Manoor P. Hande, Miquel A. Pujana, Razqallah Hakem, Karim Mekhail

## Abstract

The nuclear envelope is a membrane separating nuclear from cytoplasmic processes. Existing models suggest that damaged DNA moves to the envelope at the edge of the nucleus for repair. Yet, most damaged human DNA does not reposition to the nuclear periphery during repair. Here we show that human cells relocate the nuclear envelope to non-peripheral damaged DNA, providing solid support promoting the reconnection of DNA break ends. Upon DNA double-strand break (DSB) induction, cytoplasmic microtubules poke the nuclear envelope inwards, inducing an extensive network of DSB-capturing nuclear envelope tubules (dsbNETs). The formation of dsbNETs, which encompass the nuclear lamina and the inner and outer nuclear membranes, depends on DNA damage response kinases, dynamic microtubules, the linker of the nucleoskeleton and cytoskeleton (LINC) proteins SUN1 and SUN2, nuclear pore protein NUP153, and kinesin KIF5B. Repressing dsbNETs compromises the reassociation of DSB ends. The timely reversal of dsbNETs by the kinesin KIFC3 also promotes repair. DSB ends reconnection is restored in dsbNETs-deficient cells by enlarging the 53BP1 DNA repair center. The lamina-binding domain of SUN1 mediates its entry into the tubules and DSB capture by the envelope. Fusing truncated SUN1 to the NHEJ repair protein KU70 fails to localize SUN1 to the tubules but rescues DSB targeting only to the boundary envelope. Although dsbNETs typically promote accurate DSB repair and cell survival, they are co-opted by the PARP inhibitor olaparib to induce aberrant chromosomes restraining BRCA1-deficient breast cancer cells. We uncover dsbNETs, which bring the nuclear envelope to DSBs for repair and potentiate the efficacy of anti-cancer agents. Our findings revise theories of the structure-function relationship of the nuclear envelope and identify dsbNETs as a critical factor in DNA repair and nuclear organization, with implications for health and disease.

---

Normal and cancer cells limit excessive DNA damage to support survival and growth^1^. DNA double-strand breaks (DSBs), highly toxic lesions commonly induced by radiation or chemotherapeutics, can instigate aberrant chromosome rearrangements and stall cell growth^1–5^. Most DSBs are repaired by the primary DNA repair pathways of non-homologous end-joining (NHEJ) and homologous recombination (HR)^4–6^. NHEJ operates throughout the G_1_ and S/G_2_ stages of the cell cycle while HR is active during the S/G_2_ stage. Our understanding of NHEJ and HR has benefited from their evolutionary conservation, which facilitated their study using different model organisms. Studies in yeast and fly cells revealed that DNA damage increases the mobility of DNA^7–13^. Increased mobility facilitates temporary DSB relocation onto intranuclear microtubule or actin filaments to the nuclear periphery^11–18^. There, DSBs interact with proteins embedded within the nuclear envelope’s inner or outer nuclear membrane (INM, ONM).

Such perinuclear factors, which include the nuclear pore complex (NPC) and the linker of nucleoskeleton and cytoskeleton (LINC) complex, can facilitate NHEJ or HR via an unclear mechanism^11–20^. The LINC complex encompasses the INM-embedded SUN domain proteins SUN1 and SUN2 and the ONM-embedded KASH domain proteins called SYNEs or Nesprins. Notably, disruption of the murine or human LINC complex or its associated nuclear lamina compromises DSB mobility and repair^21–23^. Yet, most mammalian DSBs exhibit only small-scale mobility and primarily reside far from the nuclear periphery during repair^24–26^. It is unclear how the nuclear envelope promotes DNA repair in mammalian nuclei or mechanistically drives repair in any organism.

## Microtubule-dependent lamina tubules at DSBs

To study the nuclear envelope during DNA repair, we first induced DSBs using the topoisomerase II inhibitor chemotherapeutic etoposide^27^. We first visualized the nuclear envelope-associated lamina via Lamin-B1 (LMNB1) in U2OS osteosarcoma cells using immunofluorescence (IF), three-dimensional super-resolution microscopy, and in silico reconstruction of volume surfaces. Within 0.5 or 1 h of etoposide treatment, LMNB1-marked tubules emerged from around the nucleus infiltrating it (Fig. 1a-c; Supplementary Fig. 1a-d; Supplementary Movies 1 and 2). Removing etoposide for 2 h or pre-treating cells with the microtubule dynamics inhibitor nocodazole for 1 h repressed etoposide-induced LMNB1 tubules without significantly altering the distribution of cells in different cell cycle stages (Fig. 1a-c; Supplementary Fig. 1e). Etoposide induced LMNB1 tubules during both the G_1_ and S/G_2_ phase of the cell cycle (Fig. 1d). The LMNB1 tubules surrounded cytoplasmic microtubules and displaced chromatin in live and fixed cells (Fig. 1e; Supplementary Fig. 1f,g)^28^. Similar to nocodazole, treating cells with the microtubule dynamics inhibitor vinblastine prevented etoposide from inducing the LMNB1 tubules (Supplementary Fig. 1h). The LMNB1 tubules colocalized with DSBs marked by the NHEJ-promoting repair protein TP53-binding protein 1 (53BP1) (Fig. 1f-h; Supplementary Fig. 1i; Supplementary Movies 1 to 2). Consistent with these data, co-immunoprecipitation revealed that LMNB1 pulled down 53BP1 following etoposide treatment, revealing increased interaction between the endogenous proteins (Supplementary Fig. 1j). In addition, etoposide removal resulted in the reversal of the tubules, coinciding with a decrease in the number of 53BP1 DSB foci, suggesting the coordination of DSB and tubule levels (Fig. 1f-h). During time-lapse microscopy, even in the presence of etoposide, we observed the natural reversal of etoposide-induced tubules paralleled by the resealing of chromatin (Supplementary Fig. 1k; Supplementary Movies 3 and 4). Also, repressing tubule formation via pre-treatment with the microtubule dynamics inhibitors nocodazole or vinblastine exacerbated 53BP1 DSB foci accumulation upon etoposide treatment (Fig. 1f-h; Supplementary Fig. 1l). Etoposide still induced tubules coinciding with 53BP1 foci in cells treated with nocodazole as long as the latter drug is removed, and cells are allowed to recover for 1 h before retreating with etoposide (Fig. 1a-c,f-h; Supplementary Movie 2). Notably, etoposide-induced DSBs marked by the RAD51 HR protein also displayed an ability to co-localize with the LMNB1 tubules (Supplementary Fig. 1m). Considering the association of LMNB1 tubules with 53BP1 and RAD51 (Supplementary Fig. 1i,m) and the induction of the tubules in the G_1_ and S/G_2_ stages of the cell cycle (Fig. 1d), we next assessed the impact of inhibiting the DNA damage response (DDR) signaling kinases on LMNB1 tubules. The latter are the NHEJ-related DNAPK and ATM and the HR-related ATR. Also, in addition to their well-characterized role in regulating DNA repair, each of these signaling kinases is linked to the nuclear envelope proteins or microtubule-associated kinesins (Supplementary Fig. 1n). Consistent with these observations, inhibition of DNAPK, ATM, or ATR decreased the levels of etoposide-induced LMNB1 (Fig. 1i). In the absence of etoposide, the inhibition of DNAPK or ATR but not ATM repressed the baseline levels of the tubules (Fig. 1i). That the inhibition of DDR signaling kinases can decrease baseline tubules in the absence of etoposide suggests that these tubules are induced by endogenous DNA damage. Consistent with this rationale, promoting the accumulation of endogenous DSBs by knocking down the DNA repair factors KU70, RAD50, or NBS1 was sufficient to induce LMNB1 tubules (Supplementary Fig. 1o-r). Thus, endogenous or exogenous sources of DNA damage trigger transient LMNB1 tubules at different cell cycle stages. DNAPK, ATM and ATR induce LMNB1 tubules in response to exogenous DNA damage, while DNAPK and ATR but not ATM induce the tubules in the presence of endogenous DNA lesions. The tubules infiltrate chromatin and capture DSBs marked by DNA repair proteins promoting NHEJ or HR repair.

**Fig. 1.**
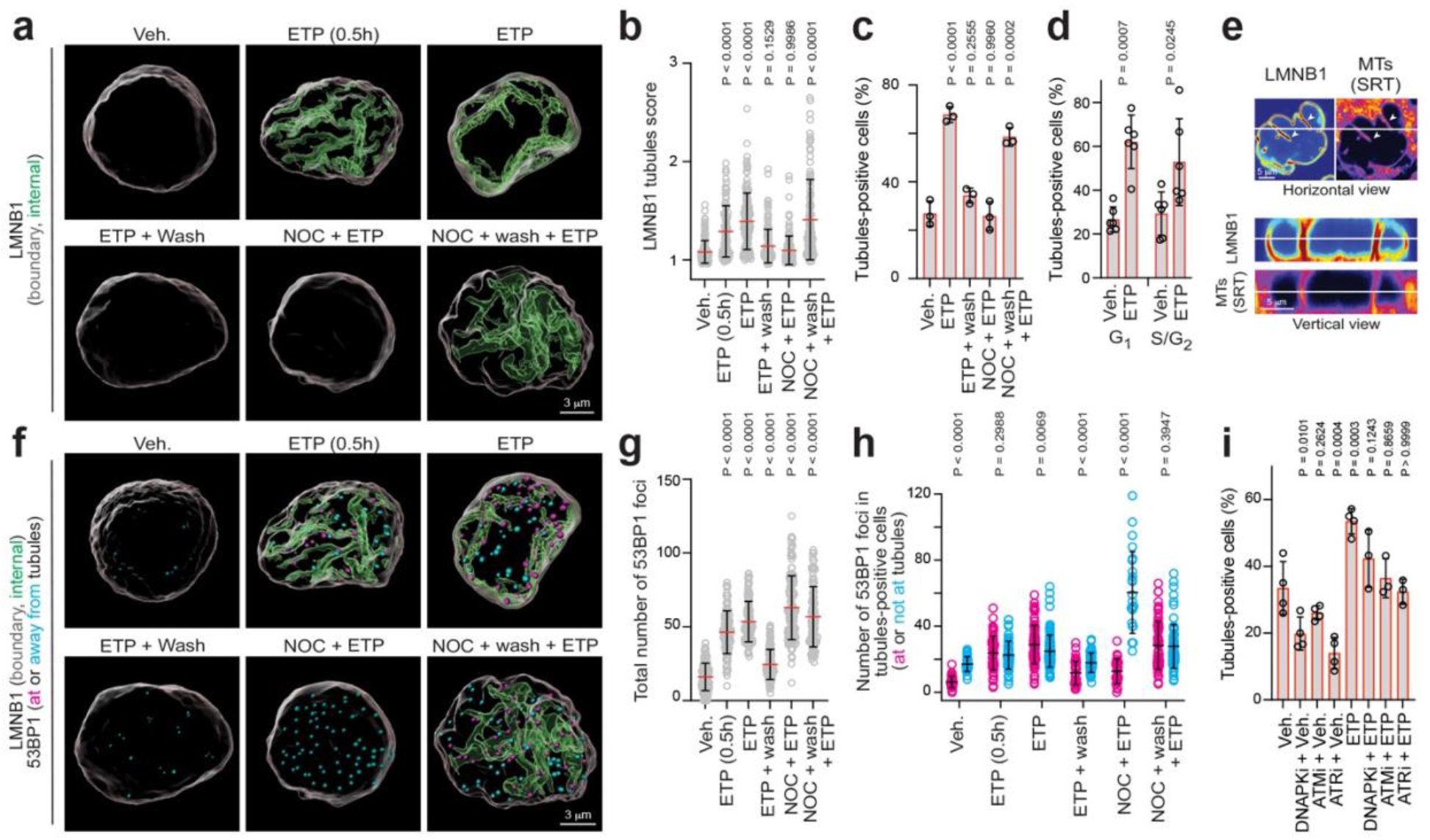
DSBs at LMNB1 tubules driven by cytoplasmic microtubules and the DNA damage response. (a-c) Representative images (a) and quantifications (b-c) showing etoposide (ETP)-induced intranuclear LMNB1 tubules and their reversal upon removing the drug (wash) or their prevention by nocodazole (NOC). (d) Etoposide-induced LMNB1 tubules during the G_1_ and S/G_2_ cell cycle stages. (e) Etoposide-induced LMNB1 tubules infiltrated by cytoplasmic microtubules in live cells. (f-h) Etoposide-induced LMNB1 tubules and 53BP1-marked DSBs. Representative images (f), 53BP1 nuclear foci counts presented as total (g) or according to their position at (magenta) or away from (cyan) the LMNB1 tubules (h). (i) Effect of treatment with different DDR signalling kinase inhibitors (DNAPKi, ATMi, and ATRi) on LMNB1 tubule formation. (a-i) U2OS cells; etoposide (ETP) was 100 μM for 1h unless otherwise indicated; data are shown as the mean±SD; n=3 (b,c,g,h), n=6 (d), or n=4 or n=3 (i) biologically independent replicates; two-way ANOVA with Dunnett’s (b,c,h,i) or Tukey’s (d) multiple comparisons tests, and multiple unpaired t-tests (g).

The stronger induction of LMNB1 tubules in the G_1_ phase compared to the S/G_2_ stages of the cell cycle (Fig. 1d) may be related to a small but significant bias for NHEJ and HR repair to occur within the central and peripheral zones of the nucleus, respectively (Supplementary Fig. 2a). This observation is consistent with the higher reliance of perinuclear heterochromatin on HR-based repair^7,8,14–17,29–31^. DSBs and LMNB1 tubules were efficiently induced at all etoposide concentrations tested (Supplementary Fig. 2b,c). Of note, a small percentage of etoposide-treated cells exhibited various rare phenotypes. For example, we observed severe DSB clustering in cells with no or few LMNB1 tubules and detected cells with nuclear envelope blebbing or failure (Supplementary Fig. 2d). The clustering of a high number of DSBs has been observed in yeast and human cells, particularly when lesions occur in transcriptionally active human genomic regions^15,32,33^. Cells exhibiting these rare phenotypes within our experimental conditions were excluded from analyses. Also, etoposide triggered LMNB1 tubules in different cell types, including non-cancerous cells (Supplementary Fig. 2e,f), and inhibition of transcription or the proteasome moderately increased the levels of 53BP1 foci and LMNB1 tubules (Supplementary Fig. 2g,h). So, we have observed LMNB1 tubules, which can be triggered by various DNA damage sources, in all tested cell types.

## Repair by promoters and repressors of the tubules

To determine if the LMNB1-positive tubules and their impact on DSBs are influenced by nuclear envelope-embedded proteins linked to genome stability in different organisms^26,34^, we studied the microtubule-associated LINC complex, kinesin motor proteins, and nuclear pore complex proteins^26,34^. In addition to being LMNB1-positive, the tubules harbored the INM-marking LINC subunit SUN2 protein, ONM-marking LINC subunit SYNE1 (a.k.a. Nesprin-1), and the nuclear pore complex (NPC)-indicating NUP98 (Fig. 2a). Given that the lamina, INM, and ONM were all constituents of the tubules, we dubbed them DSB-capturing nuclear envelope tubules (dsbNETs). Knockdown of the LINC subunits SUN1 or SUN2 repressed the induction of dsbNETs by etoposide and increased the accumulation of 53BP1-marked DSBs (Fig. 2b; Supplementary Fig. 3). Notably, despite the induction of 53BP1 foci upon knockdown of SUN1 or SUN2, dsbNETs formation depended more on SUN1 than SUN2, suggesting the two proteins contribute to genome stability via at least partly independent processes (Fig. 2b). In addition to the LINC complex, the nuclear pore complex is another nuclear envelope constituent that promotes genome stability via various mechanisms^34,35^. Notably, the NPC central channel protein NUP98 has been linked to actin-dependent DSB mobility and repair^30,31^ and the NPC basket component NUP153 promotes genome stability through various processes such as promoting the nuclear import of 53BP1^36–38^. We observed that NUP153 knockdown strongly repressed dsbNETs formation and induced γH2AX but not 53BP1 foci in cells treated with etoposide (Fig. 2c,d; Supplementary Fig. 3). In contrast, NUP98 knockdown induced 53BP1 foci and dsbNETs, suggesting that this protein promotes genome stability independently of the tubules (Fig. 2c). Our results reveal that SUN proteins and the NPC basket component NUP153 promote dsbNETs formation.

**Fig. 2.**
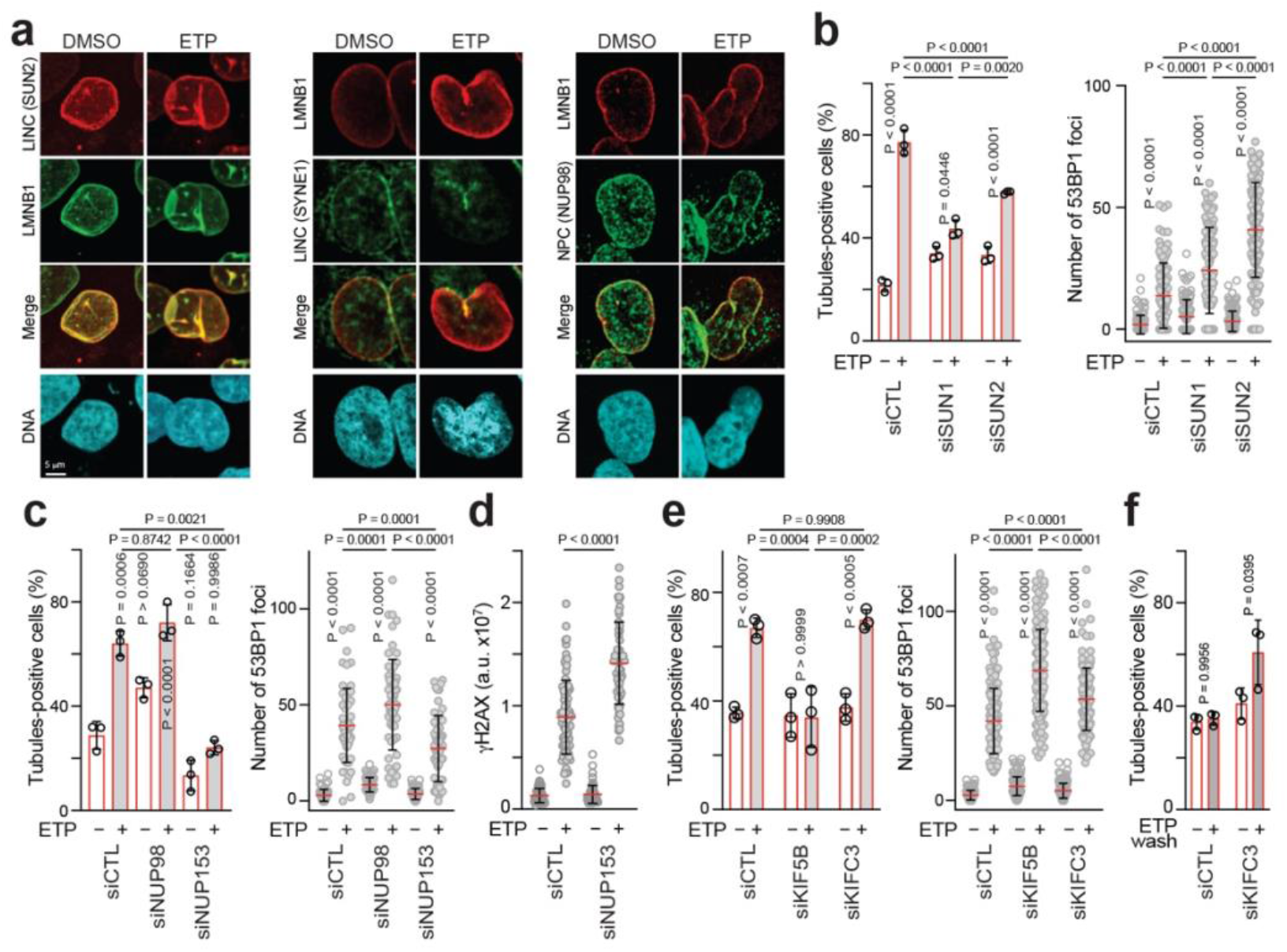
Factors controlling the formation of DNA damage-inducible LMNB1 tubules. (a) Etoposide-induced LMNB1 tubules colocalize with the LINC subunits SUN2 and Nesprin-1 and the NPC factor NUP98. (b-e) Effect of knocking down LINC proteins SUN1 or SUN2 (b), NPC subunit NUP98 or NUP153 (c,d), or the kinesins KIF5B or KIFC3 (e) on etoposide-induced LMNB1 tubule formation and the accumulation of 53BP1 or γH2AX as indicated. (f) Effect of KIFC3 knockdown on the reversal of etoposide-induced LMNB1 following removal of the drug. (a-e) U2OS cells; data are shown as the mean±SD; n=3 biologically independent replicates, two-way ANOVA with Sidak’s (b(left)) or Tukey’s (b(right) and c-f) multiple comparisons tests.

Considering the dependence of dsbNETs on microtubules (Fig. 1a-c,e; Supplementary Fig. 1f,g,h,l), the kinesin-dependent relocation of DSBs along intranuclear microtubule filaments to the nuclear periphery for repair in yeast^11,14,16^, and the dependence of mammalian DSB mobility and repair on kinesins^10,21^, we asked whether kinesins regulate dsbNETs. Cultured mammalian cells can exhibit diverse arrangements of cytoplasmic microtubules, which are generally laid with their static or minus-end at the microtubule-organizing center and their dynamic or plus-end towards the cell membrane or the ONM^12,39^. So, we considered plus-end and minus-end-directed kinesins as putative promoters and repressors of dsbNETs, excluding kinesins with critical roles in mitosis^14,21,40^. Knockdown of the microtubule plus-end-directed kinesin-1 KIF5B repressed dsbNETs and induced 53BP1 foci upon treatment with etoposide (Fig. 2e; Supplementary Fig. 3). In contrast, knockdown of the microtubule minus-end-directed kinesin-14 KIFC3 did not limit the induction of dsbNETs but still exacerbated DSB accumulation (Fig. 2e; Supplementary Fig. 3). Moreover, KIFC3 knockdown compromised the reversal of dsbNETs upon etoposide removal (Fig. 2f), consistent with this kinesin contributing to repair by ensuring the timely reversal of the tubules. Together, our data indicate that SUN1, SUN2, NUP153, and KIF5B promote the formation of transient dsbNETs that are reversed via KIFC3 to promote DNA repair.

## dsbNETs connect DSB ends

We next asked whether triggering DNA damage at a single genomic locus induces dsbNETs. We used the mCherry-tagged FokI endonuclease reporter DSB (FokI-DSB) system to trigger the lesions at a single chromosomal locus on chromosome 1 (Supplementary Fig. 4a)^41,42^. FokI-DSB induction was confirmed on a single cell level by the emergence of a mCherry-marked focus colocalizing with 53BP1 (Fig. 3a; Supplementary Fig. 4b)^41^. Cells with an induced FokI-DSB exhibited nuclear envelope tubules that were shorter and less extensive compared to those observed upon treatment with chemotherapeutic agents (Fig. 3a,b; compare to Fig. 1a and Supplementary Fig. 1a). The less extensive FokI-DSB-induced LMNB1 tubules were still repressed by the microtubule dynamics inhibitor nocodazole (Fig. 3b). Given the transient role of dsbNETs at DNA lesions, we expected it to be challenging to visualize nuclear envelope tubules interacting with the single damaged genomic locus. However, FokI-DSB co-localized with LMNB1 more at the tubules than at the boundary of the nucleus under steady-state conditions (Fig. 3a; Supplementary Fig. 4c). In addition, knockdown of SUN1, SUN2, NUP153, or KIF5B repressed FokI-DSB-induced dsbNETs and increased the distance between the DNA lesion and LMNB1 (Fig. 3c,d; Supplementary Fig. 4d-f). Moreover, knockdown of SUN1 increased the distance between the DSB and the nuclear edge at least partly independently of histone deacetylase inhibition (Supplementary Fig. 4g). This observation suggests that the impact of SUN1 on DSB positioning does not simply reflect a decrease in perinuclear silent chromatin. Our data indicate that DNA damage at a single chromosomal locus triggers dsbNETs that associate with the damaged locus.

**Fig. 3.**
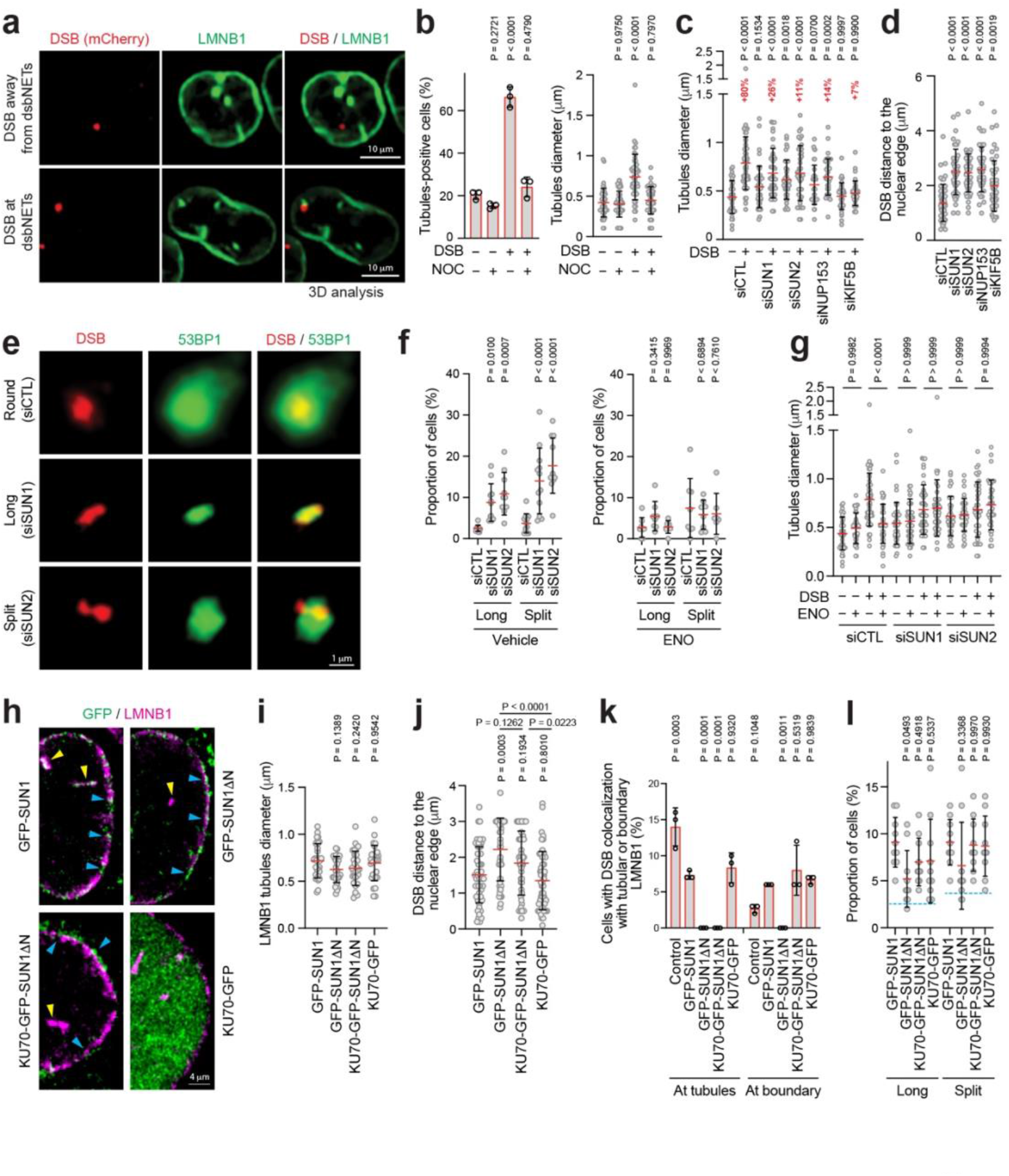
Connection of DSB ends by dsbNETs and the licensing of SUN1 for dsbNETs and DSB tethering via the nuclear lamina. (a) Association of FokI-DSB with LMNB1 tubules. (b) FokI-DSB induction triggers LMNB1 tubules as revealed by the increased percentage of cells with tubules (left) and their increased diameter (right). Tubule induction by Fok1-DSB was repressed by nocodazole (NOC). (c) Effect of disrupting LINC, NPC, or kinesin proteins on the induction of LMNB1 tubules following FokI-DSB activation. (d) Disruption of LINC, NPC, or kinesin proteins linked to dsbNET formation increases the distance between the FokI-DSB and the nuclear edge. (e-f) Effect of SUN1 or SUN2 knockdown on FokI-DSB shape. Representative images (e) and quantifications from cells treated with vehicle (f, left) or enoxacin (f, right) are shown. (g) Effect of LINC protein knockdowns and enoxacin treatment on LMNB1 tubule induction in cells without or with FokI-DSB activation. Data in (c) are reshown in (g) as they were part of the same experiment and to facilitate communication. (h) The N-terminal lamina/chromatin interaction domain of SUN1 is required for the protein to access LMNB1 tubules (yellow arrows) induced by FokI-DSB activation. The N-terminal domain is not required for SUN1 to colocalize with LMNB1 at the nuclear envelope. The N-terminal domain is present in the GFP-SUN1 but missing from the GFP-SUN1ΔN or KU70-GFP-SUN1ΔN proteins. KU70-GFP served as an additional negative control. (i) LMNB1 tubules similarly form following FokI-DSB activation in cells expressing the indicated fusion proteins. (j) Expression of GFP-SUN1ΔN but not KU70-GFP-SUN1ΔN increases the distance between the FokI-DSB and the nuclear edge. (k) Assessment of the colocalization of FokI-DSB with tubular or boundary LMNB1 in cells expressing the indicated chimeric proteins. (l) Effect of overexpressing fusion proteins on the shape of the DSB. Cyan lines indicate the baseline levels in wild-type cells not expressing any of the chimeric fusion proteins. (a-l) 2-6-5 FokI-DSB cells; data are shown as the mean±SD; n=3 biologically independent replicates, two-way ANOVA with Dunnett’s (b-d,f,i,j) or Tukey’s (g,k,l) multiple comparisons tests.

To gain insight into the mechanistic role of dsbNETs at the DNA lesion, we closely assessed the structure of the mCherry-marked DSB site in wild-type cells and in SUN1 or SUN2-deficient cells. Closer examination revealed that the knockdown of SUN1 or SUN2 increased the percentage of DSB foci with long or split shapes (Fig. 3e and 3f, left). The long and split mCherry-marked foci co-localized with 53BP1 foci, confirming their engagement with repair processes (Fig. 3e). The split DSB shapes had one lobe protruding from the 53BP1 repair focus (Fig. 3e). Moreover, marking one DSB end using sequence-specific short guide RNAs (sgRNAs) and catalytically dead Cas9 fused to a green fluorescent protein (dCas9-GFP) fully labeled round DSBs but mainly labeled only one side of the elongated or split DSBs (Supplementary Fig. 4h,i). This result confirms that the non-spherical DSB shapes reflect loosely connected or disconnected break ends, consistent with the gradual alignment of DSB ends during DNA damage repair^43,44^. Together, our findings suggest that dsbNETs help sequester DSB ends within the 53BP1 DNA repair focus facilitating the reconnection of DSB ends. Therefore, we envisaged that enlarging 53BP1 repair foci may restore DSB ends in LINC protein-deficient cells. Indeed, expanding the size of 53BP1 foci using the DNA repair-promoting drug enoxacin^45^ in cells deficient in SUN1 or SUN2 reconnected the DSB ends (Fig. 3e,f; Supplementary Fig. 4j-l), bypassing the need for dsbNETs (Fig. 3g). These results indicate that dsbNETs promote the reconnection of DSB ends, thereby sequestering them within the 53BP1 repair focus. Disruption of the SUN proteins compromises dsbNETs, disconnecting the DSB ends which can then escape from the DNA repair focus. Enlarging the repair focus bypasses the need for dsbNETs, reconnecting break ends in LINC-deficient cells.

## SUN1 licensing for dsbNETs and perinuclear DSB tethering

The nuclear envelope-embedded SUN1 protein relies on its N-terminal region to interact with the nuclear lamina and chromatin (Supplementary Fig. 5a)^46–50^. Therefore, we engineered and expressed an enhanced GFP-tagged version of SUN1 that was either full-length (GFP-SUN1) or lacking the lamina and chromatin interaction domain (GFP-SUN1ΔN) (Fig. 3h; Supplementary Fig. 5b,c). The truncated SUN1 protein lacks the lamina and chromatin-binding domain but retains the SUN and transmembrane domains mediating LINC assembly and association with the nuclear envelope, respectively^46–50^. We included an additional chimeric protein in which we fused GFP-SUN1ΔN to the KU70 NHEJ protein, which is not required for tubule formation (Supplementary Fig. 1o), creating KU70-GFP-SUN1ΔN. We also used KU70-GFP alone as an additional control. Although we engineered all SUN1 fusion proteins to be resistant to a siRNA targeting endogenous SUN1 (Supplementary Fig. 5b,c), expression of the chimeric full-length or truncated SUN1 fusion proteins was well tolerated by FokI-DSB cells only in the presence of endogenous SUN1. Therefore, we assessed the effect of expressing the chimeric proteins in FokI-DSB cells harboring endogenous SUN1. The GFP-SUN1, GFP-SUN1ΔN, or KU70-GFP-SUN1ΔN proteins displayed a similar localization pattern surrounding LMNB1 signal at the edge of the nucleus while KU70-GFP was nucleoplasmic, as expected (Fig. 3h). Although LMNB1-marked tubules similarly formed upon expressing each of the chimeric proteins, GFP-SUN1 but not GFP-SUN1ΔN or KU70-GFP-SUN1ΔN entered the LMNB1-marked tubules (Fig. 3h,i). This observation is consistent with the LMNB1 tubules being dependent on endogenous SUN1 (Fig. 3c) and indicates that the chromatin/lamina-interacting domain of the GFP-tagged SUN1 protein either provides it with access to or retains it within the LMNB1-marked tubules.

Compared to GFP-SUN1, expression of GFP-SUN1ΔN but not KU70-GFP-SUN1ΔN increased the distance between the DSB and nuclear edge (Fig. 3j). Also compared to GFP-SUN1, GFP-SUN1ΔN decreased the colocalization of the DSB with LMNB1 both at the tubules and the nuclear boundary (Fig. 3k). Notably, fusing GFP-SUN1ΔN to KU70 restored the colocalization of the DSB with LMNB1 at the nuclear boundary but not the tubules (Fig. 3k). In addition, cells expressing GFP-SUN1, GFP-SUN1ΔN, or KU70-GFP-SUN1ΔN all showed elevated levels of long and split DSB shapes (Fig. 3l), suggesting that overexpression of any SUN1-containing chimeric protein limits the ability of SUN1 to connect DSB ends to each other. Taken together, these findings indicate that the N-terminal domain of SUN1 allows it to stably localize to dsbNETs and is required for the protein’s ability to capture DSBs (Supplementary Fig. 5d). Also, the results reveal that fusing the truncated SUN1 protein to KU70 restored the protein’s ability to capture DSBs at the nuclear boundary (Supplementary Fig. 5d).

Of note, immunoblotting revealed that SUN1 migrates to a few differently sized bands repressible by SUN1 knockdown (Supplementary Fig. 5e). Also, etoposide altered the relative abundance of the different SUN1 bands suggesting a complex regulation of the protein during DNA repair (Supplementary Fig. 5f). In contrast, etoposide treatment lowered SUN2 expression (Supplementary Fig. 5g), pointing to a potential stoichiometric reconfiguration of SUN1 and SUN2-containing complexes during DNA damage repair. Consistent with these data, etoposide treatment increased the general protein phosphoserine signal co-immunoprecipitating with SUN1 (Supplementary Fig. 5h). In contrast, etoposide treatment did not alter the protein ubiquitylation signals in SUN1 pulldowns (Supplementary Fig. 5h,i). Our data suggest that SUN1 or its binding partners may be subject to serine phosphorylation as part of a complex reconfiguration of LINC proteins during DNA repair.

## Targeting cancer via dsbNETs

Considering the connections between dsbNETs and DSBs and the generally increased levels of DNA damage in cancer, we next explored the potential impact of dsbNETs on different cancers. Given that KIF5B induces while KIFC3 reverses dsbNETs, we analyzed cancer RNA-Seq datasets (TCGA, TARGET, and ICGC; see methods)^51^ to determine significant correlations between the expression of *KIF5B* or *KIFC3* and the expression of various DNA repair factors. Hierarchal clustering revealed that the expression of *KIF5B* and *KIFC3* positively and negatively correlated with the expression of DNA repair proteins involved in different repair pathways across cancers, respectively (Supplementary Fig. 6a,b). *KIF5B* expression positively correlated with an NHEJ gene expression signature in 92% (23/25) of the analyzed cancer types, while *KIFC3* expression negatively correlated with the same signature in 84% (21/25) of the cancers (Fig. 4a; Supplementary Fig. 6c). The bias towards a positive correlation of *KIF5B* and negative correlation of *KIFC3* with the NHEJ signature was much stronger than with signatures representing the HR or alternative end-joining (altEJ) pathways of DNA repair (Fig. 4a,b; Supplementary Fig. 6c-e). The bias towards a positive correlation of *KIF5B* and negative correlation of *KIFC3* was observed for breast cancer for both of the NHEJ and HR signatures, albeit the effect was more pronounced for the NHEJ signature (Fig. 4a,b). These results suggest that by coordinating the expression of bone fide DNA repair factors and dsbNET regulators, cancer cells may better tolerate their relatively higher DNA damage load. Consistent with these findings, human breast tumor sections with mutations in the HR-mediating breast and ovarian cancer genes *BRCA1* or *BRCA2* showed nuclei with nuclear membrane grooves (Supplementary Fig. 7a,b)^51^. These features were reminiscent of the nuclear membrane grooves observed on histopathologic examination of several tumor types including Langerhans cell histiocytosis, adult granulosa cell tumor, and papillary thyroid carcinoma^52,53^. This characteristic finding can be a useful diagnostic clue for pathologists, but the pathophysiology leading to this morphologic attribute remains poorly understood. So, we further explored the possible connection between dsbNETs and breast cancer cells in cell culture.

**Fig. 4.**
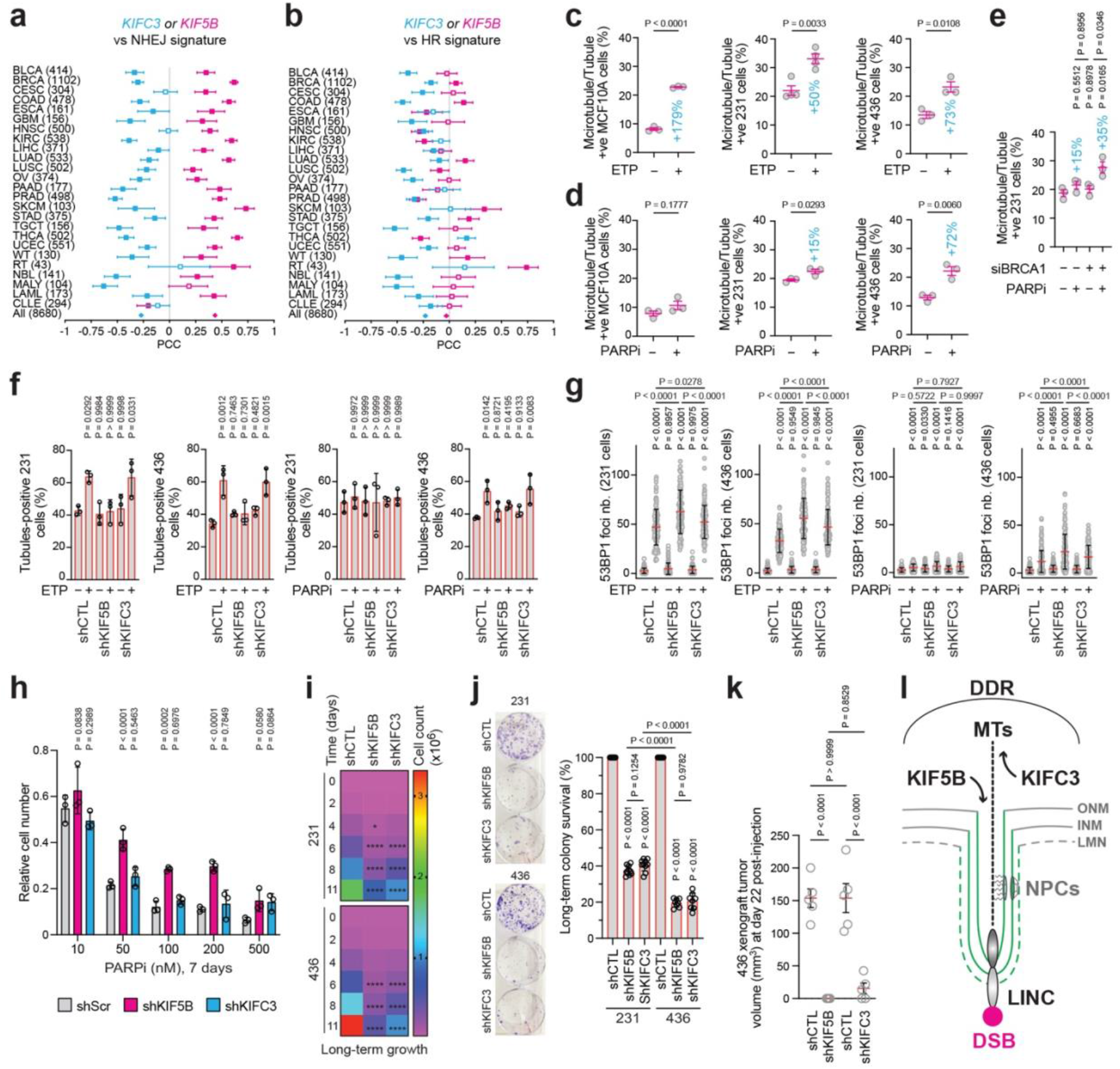
Restraining cancer cells via dsbNETs modulation. (a) The expression of *KIF5B* and *KIFC3* positively and negatively correlated with the NHEJ pathway signature across cancers, respectively. Tumor numbers are shown with cancer acronyms (BRCA, breast cancer). Significant Pearson’s correlation coefficients (PCCs) are depicted for *KIF5B* and *KIFC3* in magenta and cyan-filled data points, respectively. White squares are non-significant. (b) Same as in (a) but shows the PCCs for the HR signature. (c) Etoposide treatment (ETP, 100 μM, 1h) induced microtubule-positive LMNB1 tubules in the non-tumorigenic MCF10A breast cells, BRCA1-proficient TNBC cells (MDA-MB-231; 231), and BRCA1-deficient TNBC cells (MDA-MB-436; 436). (d) The PARP inhibitor olaparib (PARPi, 2 μM, 1 h) induced microtubule-positive LMNB1 tubules more in MDA-MB-436 cells than in MDA-MB-231 or MCF10A cells. (e) BRCA1 knockdown promoted the formation of microtubule-positive tubules in MDA-MB-231 cells treated with PARPi (PARPi, 2 μM, 1 h). (f,g) Effects of KIF5B or KIFC3 knockdown on LMNB1 tubules (f) and 53BP1 foci number (g) in cells treated with etoposide (ETP, 100 μM, 1 h) or olaparib (PARPi, 2 μM, 24 h). (h) Knockdown of KIF5B, but not KIFC3, provides resistance to the PARPi olaparib in a long-term growth assay. (i,j) Knockdown of KIF5B or KIFC3 decreased relative long-term cell growth in MDA-MB-231 and especially in MDA-MB-436 cells as assessed by standard cell culture (i) and colony formation assays (j) in the absence of DNA-damaging agents. (k) Murine xenografts showing that knockdown of KIF5B or KIFC3 is sufficient to restrain the ability of MDA-MB-436 cells to form tumors in murine xenografts 22 days-post injection. (l) General dsbNETs model. See the main text or Supplementary Fig. 8 for details. DDR, DNA damage response signaling kinases ATR, DNAPK, and ATM; MTs, microtubules; KIF5B, MT plus-end-directed kinesin-1; KIFC3, MT minus-end-directed kinesin-14; NPC, nuclear pore complex; LINC, linker of nucleoskeleton and cytoskeleton; DSB, DNA double-strand break. (c-k) Data are shown as mean±SD, n=3 biologically independent replicates for all quantifications, except n=5 mice per condition (k); two-tailed t-test (c-e), two-way ANOVA with Tukey’s multiple comparisons test (f-h) or Dunnett’s multiple comparisons test (i), and one-way ANOVA with Tukey’s multiple comparisons test (k).

We first used aggressive triple-negative breast cancer (TNBC) cells with wild-type (MDA-MB-231) or mutant (MDA-MB-436) *BRCA1* as well as the non-tumorigenic MCF10A mammary epithelial cells. We detected microtubule-positive LMNB1 tubules in all cell types, especially upon treatment with etoposide (Fig. 4c). *BRCA1*-mutant cells lack HR, over-rely on NHEJ and other repair pathways, and are preferentially killed by the clinically approved PARP inhibitor olaparib, which triggers deleterious chromosome structures such as radials^3,5,21,42,54–58^. Unlike etoposide treatment (Fig. 4c), the relative induction of microtubule-positive LMNB1 tubules by olaparib was more pronounced in *BRCA1*-mutant TNBC cells compared to *BRCA1*-proficient TNBC or MCF10A cells (Fig. 4d). Moreover, transient knockdown of BRCA1 in MDA-MB-231 cells increased tubule formation upon olaparib treatment (Fig. 4e; Supplementary Fig. 7c). These results indicate that BRCA1 deficiency exacerbates the induction of dsbNETs in TNBC cells upon exposure to the PARPi olaparib as compared to the topoisomerase II inhibitor etoposide.

To assess the impact of dsbNETs on breast cancer cells, we knocked down KIF5B or KIFC3 using a doxycycline-inducible short hairpin RNA (shKIF5B or shKIFC3) in MDA-MB-231 and MDA-MB-436 cells (Supplementary Fig. 7d,e). Knockdown of KIF5B but not KIFC3 decreased the levels of dsbNETs induced by etoposide or olaparib (Fig. 4f), as expected (Fig. 2e, left). Also, depletion of KIF5B or KIFC3 increased etoposide and olaparib-induced DSB levels in short-term cell cultures (Fig. 4g). Notably, *BRCA1*-mutant breast cancer cells are sensitive to PARPi, but the development of resistance to this life-saving class of drugs can underly clinical relapse^5,42,59–62^. So, there exists a need to identify mechanisms that potentiate cellular sensitivity to PAPRi. Notably, disruption of KIF5B but not KIFC3 decreased the sensitivity of MDA-MB-436 cells to PARPi in long-term cultures (Fig. 4h; 7 days). Similarly, the ability of PARPi to eventually induce the formation of toxic radial chromosomes in MDA-MB-436 cells decreased following KIF5B knockdown (−18.2%) and slightly increased following KIFC3 knockdown (+5.1%) (Supplementary Fig. 7f). Together, our results suggest that the transient formation of dsbNETs within 24 h of DNA damage induction lowers DSB levels in cells treated with etoposide or PARPi, likely through the quick action of NHEJ. However, the ability of PARPi to induce the highly toxic radial chromosomes in HR/BRCA1-deficient cells likely relies on KIF5B and dsbNETs. Consistent with this notion, an elegant study previously revealed that the ability of PARPi to induce NHEJ-dependent radials in BRCA1-deficient murine cells was mediated by SUN1, SUN2, and dynamic microtubules^12,21^. This is also consistent with the role of the nuclear envelope proteins and kinesins in triggering deleterious break-induced replication repair at subtelomeric DSBs in budding yeast^14^.

We noted that depleting KIF5B or KIFC3 in the absence of olaparib still induced relatively low levels of radials in *BRCA1*-mutant TNBC cells (Supplementary Fig. 7f). So, we assessed the impact of depleting the kinesins on long-term cancer cell growth in the absence of exogenous DNA damaging agents. Indeed, knockdown of KIF5B or KIFC3 did not alter cell growth in the short-term (2-4 days) but eventually lowered growth (6-11 days), especially in *BRCA1*-mutant MDA-MB-436 cells (Fig. 4i). Assays assessing the long-term ability of cells to grow into colonies yielded similar results (Fig. 4j). In addition, knockdown of KIF5B or KIFC3 decreased the ability of MDA-MB-436 TNBC cells to form tumors in mouse xenografts (Fig. 4k). Consistent with our findings, reanalysis of genomic CRISPR screen data from DepMap^63^ further confirmed a synthetic lethal interaction between BRCA1 and KIF5B (Supplementary Fig. 7g). Together with prior studies^12,14,21^, our data indicate that disrupting dsbNETs alone decreases faithful repair and induces ectopic repair that eventually hinders cell growth. However, in the context of BRCA1 deficiency, the ability of PARPi to induce cancer cell-killing aberrant chromosome structures via NHEJ partly depends on dsbNET-inducing factors, including KIF5B, SUN1/2, and dynamic microtubules.

## Discussion

Our results identify dsbNETs and their regulators and roles in promoting faithful DNA damage repair and cell growth and survival (Fig. 4l; Supplementary Fig. 8). Upon DSB induction, DDR signaling cooperates with microtubules to induce dsbNETs, which then infiltrate the nucleus. The dsbNETs encompass the INM, ONM, and nuclear lamina and are promoted by the DDR kinases, LINC proteins SUN1 and SUN2, the NPC basket component NUP153, and the kinesin-1 KIF5B (Supplementary Fig. 8a). This internal nuclear envelope network composed of dsbNETs captures DSBs, providing the solid support necessary to efficiently reconnect the ends of a DSB to each other, ensuring accurate DNA repair. Disrupting dsbNETs weakens the connection between the DSB ends, which can then escape the 53BP1 DNA repair center. Enlarging 53BP1 foci restores the localization of DSB ends within DNA repair foci, bypassing dsbNETs and re-reconnecting DSB ends. The lamina and chromatin interaction domain of SUN1 licenses it for entry into the dsbNETs and mediates DSB tethering to the nuclear envelope. Under standard conditions, disrupting dsbNETs increases the chance of mis-joining the ends of DSBs on different chromosomes, mediating misrepair and hindering growth (Supplementary Fig. 8b). In the context of BRCA1-deficient breast cancer cells treated with PARPi, the ability of the drug to induce cancer cell-killing aberrant chromosome structures is boosted by dsbNETs (Supplementary Fig. 8c).

Our work reveals a unifying model that explains how DSB-nuclear envelope interactions promote DNA repair and genome stability in different organisms. In yeast and fly cells, the higher degree and scale of DSB mobility allows the DNA lesions to diffuse or be transported onto nuclear filaments to the nuclear envelope at the nuclear edge for repair^7–9,16,30,64–67^. Here we uncover that human cells deploy dsbNETs throughout the nucleus, overcoming the relatively low mobility of human DSBs^10,24,25,68^. So, dsbNETs can readily bring the envelope to the DSBs in mammals instead of vice versa, as in lower eukaryotes. Yet, we must highlight that many DSBs and telomeres undergoing telomerase-independent alternative lengthening can exhibit long-range mobility in mammalian nuclei^33,69,70^. Also, we only captured about half of the studied DSBs associating with dsbNETs. So, it remains possible that a sizable proportion of human DSBs may be targeted to the nuclear edge or even repaired without contacting the envelope. Consistent with this notion, kinesin and myosin motor proteins as well as microtubule and actin filaments promote DSB mobility and repair in yeast, fly, murine or human cells^14,21,30,31,71–75^. Thus, together with the existing literature, our findings reveal that DSBs and the nuclear envelope have likely been connecting for billions of years by employing similar molecules. Yet, each organism appears to organize the conserved molecules in unique fashions best suited to the specific characteristics and challenges faced by a given genome in nuclear space. The herein-uncovered convergence of these independent lines of research employing different genetic models further emphasizes the seminal contributions of genetic model organisms to our understanding of genome organization and stability.

The nuclear envelope likely provides chromosomes with diverse types of solid support to ensure chromosome stability. Here we uncover that the envelope promotes the faithful reconnection of human DSB ends during DNA repair. This is reminiscent of the solid support provided by the yeast nuclear envelope to tether sister chromatids to each other during replication and prevent lifespan-shortening unequal sister chromatid exchanges within DNA repeats at the nuclear edge^29,34,76,77^. This mechanism promotes replicative lifespan in yeast by maintaining ribosomal DNA (rDNA) and telomeric stability^29^. Further supporting the possibility that the human nuclear envelope may provide solid support to rDNA repeats, damaged rDNA repeats relocate from the center to the periphery of the nucleolus and interact with SUN1 proteins in human cells^78,79^. Notably, mobility of the DSB-like uncapped telomeres and telomere-telomere fusions are regulated by microtubules and kinesins, including KIF5B, in murine cells^21^. We note that dsbNETs or similar structures may help repair distinct types of DNA lesions or provide solid support to various molecular pathways.

Nuclear envelope grooves have been especially observed in different pathological settings^52,53^. Indeed, current models suggest that the interphase nuclear envelope in healthy cells is a relatively smooth structure that becomes severely deformed in settings such as aggressive cancers, premature aging, and advanced natural aging^22,23,34,76,80–88^. In contrast, our results indicate that the interphase nuclear envelope structure can rapidly and drastically transform to exert functions critical to genome stability and cell survival. This raises a number of key points that will require further consideration by different fields intersecting at the levels of nuclear structure, genome organization and stability. Taken together, our study redefines theories of the structure-function relationship of the human nuclear envelope and invites us to reassess our understanding of the nucleus in different health and disease settings.

## Acknowledgments

We thank Dr. Kimberly Lau and Paul Paroutis at the SickKids Children’s Hospital Imaging Facility, Dr. Feng Xu at the Advanced Optical Microscopy Facility of the Princess Margaret Cancer Research Tower, and the Imaris Bitplane support team. We thank Roger A. Greenberg for sharing the FokI-DSB system and Shane Harding for the NBS1 antibody. We thank members of the labs of K. Mekhail, R. Hakem, and M. Ohh for their support and feedback. We thank Daniel Durocher and Irene Chiolo for their fruitful feedback. The images and quantification of nuclear groves from representative cases of human breast carcinoma were obtained from data generated by the TCGA Research Network (https://www.cancer.gov/tcga). M. Shokrollahi is supported by the Ontario Graduate Scholarship program. A. Hundal is supported by the Ontario Women’s Health Scholars Award and the Canadian Institutes of Health Research (CIHR) scholarship program. P.G. Maass was supported by funds from the CIHR (173542). M.A. Pujana was supported by funds from Instituto Salud Carlos III (PI21/01306 and CIBERES, co-funded by the European Regional Development Fund ERDF, a way to build Europe), Generalitat de Catalunya (SGR 2017-449; PERIS PFI-Salut SLT017-20-000076 to R Espin), and CERCA Program to IDIBELL. R. Hakem was supported by funds from CIHR (FDN143214, PJT 183597) and holds the Lau Chair in Breast Cancer Research at the University of Toronto and Princess Margaret Hospital. This research was primarily supported by CIHR funds (399687, 469861) to K. Mekhail, who also acknowledges support from the Royal Society of Canada.

## Author contributions

M.Sh., M.St., and K.M. conceived the study. K.M. together with R.H. conceptualized breast cancer studies. K.M. wrote the main text and edited the manuscript. All authors contributed to experimental design, writing, and editing of other parts of the study. M.Sh. designed and conducted the 2D and 3D for imaging, reconstruction, and related imaging, analyses, and movies. M.St. designed and conducted 2D imaging and analyses, concentric nuclear zone studies, targeted DSB and cell cycle imaging and related analyses. M.Sh., M.St., A.Hu., and J.N.Y.C performed knockdowns and related imaging and immunoblotting. A.Hu. designed and conducted coIPs, 2D imaging and analyses, metaphase spreads, and all growth assays with contributions from M.P.H to metaphase spread analyses, from A.Ha. to colony-forming assays, and from J.H to xenografts. J.N.Y.C. conducted coIPs and also designed, engineered, and validated the SUN1 chimeras, which were imaged and analyzed by M.St. The bioinformatic analyses were performed by R.E.G. and M.A.P. while J.H. analyzed CRISPR screen datasets. P.G.M. provided critical 3D imaging tools and resources. B.D. retrieved and analyzed the human tumor images. K.M. supervised M.Sh., M.St., A.Hu, J.N.Y.C., and D.U. and coordinated the project. A.Hu, A.Ha, and J.H. were supervised by R.H. while M.A.P. supervised R.E.G.

## Additional information

**Supplementary information** is available with this paper

## METHODS

### Cell culture, drug treatments, and related reagents

U2OS, IMR-90, HEK293T, and HeLa cells (cat# CCL-2, ATCC) were cultured in Dulbecco’s modified Eagle medium (DMEM, Wisent Bioproducts) supplemented with 10% fetal bovine serum (FBS, Wisent Bioproducts) and 1% penicillin/streptomycin. MDA-MB-231 and MDA-MB-436 were cultured in DMEM supplemented with 10% fetal bovine serum and 1% penicillin/streptomycin (Wisent Bioproducts). MCF10A cells were cultured in DMEM/F12 media supplemented with 17% FBS, 100 µg/mL EGF, 1 mg/mL hydrocortisone, 1 mg/mL cholera toxin, 10 mg/mL insulin, and 1% penicillin/streptomycin. The U2OS-derived ER-mCherry-LacI-FokI-DD (FokI-DSB) 2-6-5 cell line^41^ was a kind gift from Roger A. Greenberg (University of Pennsylvania). The 2-6-5 cells were grown in DMEM supplemented with 10% FBS, 1% penicillin/streptomycin, 2 μg/mL puromycin, and 200 μg/mL hygromycin. All cell lines were frequently tested for mycoplasma using MycoAlert Plus Mycoplasma Detection Kit (cat# LT07-218, Lonza).

Etoposide (cat# E7657, LKT laboratories) was added at indicated concentrations for 1h, and cells were incubated until harvesting. Wash experiments consisted of replacing the etoposide-containing media with fresh drug-free media for 2 h. Nocodazole (cat# 487928-10MG, Sigma, cat#1228 Tocris) was added at 50 μM for 1h alone or 1h prior to the addition of etoposide or DMSO vehicle control. Vinblastine (cat# V1377-10MG, Sigma-Aldrich) was added at 100 nM. DNA-PK inhibition was achieved by adding NU7026 (cat# 2828, Tocirs) at 10 μM, PARP inhibitor olaparib (AZD2281, cat# S1060, Selleckchem) was added at 1 μM, ATR inhibitor berzosertib (VE-822, cat# S7102, Selleckchem) was added at 5 μM, and ATM inhibitor (AZD1390, cat# S8680, Selleckchem) was added at 5 μM. The histone deacetylase inhibitor SAHA (Vorinostat, cat#SML0061-MG5, Millipore Sigma) was used at 10μM for 24 h.

FokI-DSB was induced in 2-6-5 cells via a 1 h treatment with 1 μM Shield-1 ligand (cat# AOB1848, AOBIOUS) and 1 μM of 4-hydroxytamoxifen (cat# S7827, Selleckchem). To enlarge 53BP1 foci, 2-6-5 cells were treated with 50 μM Enoxacin (cat# S1756, Selleckchem) for 24 h prior to DSB induction. To study LMNB1 tubules exposed to different stressors, U2OS cells were treated with 100 μM etoposide (cat# E7657, LKT laboratories) for 1 h, 50 μM enoxacin (cat# S1756, Selleckchem) for 24 h, 50 μM nocodazole (cat# 487928-10MG, Sigma) for 1 h, 2 μM flavopiridol (cat# sc-202157, Santa Cruz) for 3 h, or 10 μM MG132 (cat# 13697-25, Cayman Chemical Co) for 3 h.

### Antibodies

The primary antibodies used were PARP1 (cat# ab22724, abcam), KU70 (cat# ab83501, abcam), RAD51 (cat# ab133534, abcam), BRCA1 (cat# OP92, Millipore), gH2AX (cat# ab15083, abcam), H3ac (cat# 06-599, Millipore), SUN1 (cat# 24568-I-AP, Proteintech), SUN2 (cat# MABT880, Millipore), NUP98 (cat# MA5-14907, Invitrogen), NUP153 (cat# A301-788A, Bethyl Lab), 53BP1 (cat# ab36823, Abcam; cat# A300-272A, Bethyl Laboratories,), LMNB1 (cat# ab16048, abcam), LMNB1 (cat# ab16048, Abcam; cat# 66095-1-Ig,Proteintech,), KIF5B (cat# ab167429, abcam), KIFC3 (cat# ab154419, Abcam; cat# PA5-54359, Thermo Fisher), anti-β-actin (cat# AM4302 or cat# MA1-744 Thermo Fisher), RNF8 (cat# sc-271462, Santa Cruz), RAD50 (cat# NB100-1487,Novus), NBS1 (cat# A301-290A, Bethyl), Vinculin (cat# V9131, Millipore), phosphoserine (cat# AB1606, Millipore), and ubiquitin (cat# sc-8017, Santa Cruz). The Secondary antibodies used were Alexa-488 anti-mouse (cat# A11001, Invitrogen), Alexa-488 anti-rabbit (cat# A11008, Invitrogen), Alexa-568 anti-mouse (cat# A11004, Invitrogen), Alexa-568 anti-rabbit (cat# A11011, Invitrogen), Alexa-647 anti-mouse (cat# A21235, Invitrogen), anti-rabbit IgG (cat# ab171870, Abcam), HRP-conjugated anti-mouse-IgG (cat# NA931-1ML,Amersham), and anti-rabbit-IgG (cat# NA934-1ML, Amersham).

### Knockdowns and lentiviral preparation

To transiently knockdown factors of interest, cells were transfected for 72 h with 20-30 nM of corresponding siRNAs. The siRNAs (Supplementary Tables 1 and 2) were transfected using Lipofectamine RNAiMAX Transfection Reagent (ThermoFisher). For lentiviral preparation, HEK293T cells were co-transfected using GenJet™ (cat# SL00488, SignaGen Laboratories) lipofection with Tet-pLKO-puro (Addgene plasmid #21915) shRNA containing lentiviral constructs or a scrambled lentivrial construct (Addgene plasmid #162011) together with psPAX-2 (Addgene plasmid #12260) and pMD2.G (Addgene plasmid #12259). MDA-MB-231 and MDA-MB-436 cells were transduced 48-72 h post-HEK293T transfection with Tet-pLKO-puro lentiviruses containing shRNAs targeting human KIF5B (TRCN0000338580; 5’-CAACCGCAATTGGAGTTATAG-3’) or human KIFC3 (TRCN0000271534; 5’-TGAAGGCTGTGCACGAGAATC-3’). Transduced cells were maintained in 2 µg/mL puromycin and DMEM supplemented with 10% tetracycline-free FBS. shRNA knockdown was induced with 100 ng/mL doxycycline treatment for 96 hr. All knockdowns were successfully confirmed using immunoblotting, which in turn confirmed the specificity of the antibodies used.

### Cell cycle analysis

The cell cycle stage of cells was determined by exposing them for 2 h to 5-ethynyl-2′-deoxyuridine (EdU), using the Click-iT™ EdU Cell Proliferation Kit for Imaging, Alexa Fluor™ 488 dye (cat# C10337, ThermoFisher C10337) and following the detailed instructions from the manufacturer.

### Immunofluorescence (IF), super-resolution, and confocal microscopy

IF was performed as previously described^89,90^. Briefly, cells were seeded on glass coverslips, treated with the desired drug protocols, and fixed for 10 min using 4% formaldehyde or paraformaldehyde. Cells were washed in PBS and permeabilized for 10 min with 0.5% Triton X-100. Upon permeabilization, cells were washed in PBS/0.01% Tween (PBST) and kept overnight in 20% Glycerol/PBS. Cells were then washed with PBST and blocked for a minimum of 30 minutes in 4% BSA. Cells were incubated overnight at 4°C with a 1:500 dilution primary antibody in 1% BSA. The cells were then washed in PBST and incubated with a secondary antibody at a dilution of 1:1000 in 1% BSA for a minimum of 1h at room temperature. Cells were then washed in PBST, counterstained for 5 min with DAPI (D1306, Thermo Fisher), washed in PBS and mounted on glass slides with mowiol (cat# 81381, Sigma-Aldrich). Three-dimensional super-resolution imaging with 0.125 μm or 0.2 μm intervals was performed on a LSM880 with Airyscan fast detection system (Zeiss, SickKids imaging facility), and the oil immersion objective Plan-Apochromat 63×/1.4 oil DIC M27. ZEN black edition software edition 2.3 (Zeiss) was used for acquisition. All images shown were acquired in sensitivity-versus-resolution (SR) mode with 8 bits. Confocal three-dimensional imaging and z-slices of 0.25 μm, 0.3 μm, 0.5 μm or 0.5 μm step sizes was performed on a Leica TCS sp8 Lightning Confocal/STED microscope coupled to LasX software (Leica) using a 63x (1.4) oil-immersion objective (SickKids imaging facility), or an in-house Nikon Eclipse Ti2 C2+ confocal microscope coupled to NIS-Elements AR software (Nikon) and 100x (1.25) or 60x (0.85) OFN25 oil objectives.

### Live-cell treatment and confocal microscopy

Cells were cultured on Poly-D-Lysine coated 35 mm glass-bottom dishes (cat# P35G-1.5-14-C, MatTek). The SiR-Tubulin stain (Cytoskeleton CY-SC002) and NucBlue^TM^ (Hoechst 33342, cat# R37605, ThermoFischer) were used to visualize microtubules and DNA, respectively. To visualize the nuclear lamina, cells were transfected transiently with a GFP-LMNB1 plasmid, which was a kind gift from J. Lammerding (Cornell University). Super-resolution confocal imaging was performed using the Zeiss LSM880 Airyscan confocal microscope, and standard confocal imaging was performed using the Nikon A1R confocal microscope with a resonant scanner (AOMF imaging facility). Both microscopes were equipped with an incubation module at 37°C and 5% CO_2_. Airyscan data files were acquired in sensitivity-versus-resolution (SR) mode with 8 bits, 0.125 μm or 0.2 μm z-slices, and fitted zoom level with the oil immersion objective Plan-Apochromat 63x/1.4 oil DIC M27. A 32-channel gallium arsenide phosphide photomultiplier tube (GaAsP-PMT) area detector (Airyscan) collected a pinhole-plane image at every scan position. The Airyscan detector system enhances sensitivity 4–8-fold and resolution beyond the diffraction limit of light of up to 1.7-fold (∼130 nm) compared to standard confocal microscopes^91,92^. For all acquisitions, ∼5–10× less laser power was used compared to standard confocal microscopy. The ZEN black edition software edition 2.3 (Zeiss) was used for acquisition. Time-lapse live cell imaging was performed at 5 min intervals for 1 h with the same lense and 0.125 μm z-slices, SR mode, with the Zeiss LSM880 Airyscan confocal microscope. 405 nm laser was used to visualize DAPI, 488 nm was used to visualize GFP, and 561 nm laser was used to visualize mCherry.

Nikon A1R images were acquired with Plan-Apochromat, nano-crystal, 60x/1.4 NA, oil immersion lense. Two standard photomultiplier (PMT) detectors, two Gallium Arsenide Phosphide (GaAsP) high-sensitivity detectors and the MCL nano-drive allowed high quality of live cell samples. Nikon Element software was used for acquisition. 640 nm laser was used to visualize Cy5 (microtubule) signal. 405nm and 488nm lasers were used for visualizing DAPI and GFP signals, respectively.

### Image analyses and nuclear reconstruction

BitPlane Imaris 9.7 or 10.0 and ImageJ/FIJI software were used for single-cell image analysis^29,76,93^. Airyscan^91,92^ data files were first de-convoluted and processed in ZEN black (Zeiss) software. Maximum intensity projections were optimized using BitPlane Imaris 9.7 or 10.0 Image Proc function. Pixel intensity display settings were automatically or manually optimized using a thresholding method based on image quality and local contrast using BitPlane Imaris 9.7 or 10.0 Baseline Subtraction function followed by a Gaussian smoothing filter at a value of one.

For three-dimensional nuclear reconstruction and quantification of dsbNETs and DSB levels, the analyses were as follows. To quantify the degree of invagination of LaminB1 in a single nucleus, DAPI-stained DNA and LMNB1 signals were reconstructed in three dimensions visualizing the invaginated signal and the boundary signal separately using the Surfaces MatLAB Xtension and masking function where every z-plane in a single nucleus (30-85 z-slices) was contoured manually to ensure optimal separation of boundary and internal LMNB1 signal to detect LMNB1 nuclear tubules. For each three-dimensionally reconstructed surface, the surface area was quantified using the BitPlane Imaris 9.7 detailed Statistics function. The tubular score or invagination ratio was then quantified as the total LMNB1 surface area divided by the total DNA surface area marked by DAPI. Cells were considered tubules-positive when two or more tubules extending deeper than the radius of the nucleus were detected using BitPlane Imaris 9.7 Section function. Alternatively, z-stacks of stained cells were analyzed using the Orthogonal Views function on ImageJ/FIJI. Using BitPlane Imaris 9.7 Section, cells were considered positive for nucleus-reshaping microtubules if the latter were detected at the mid-plane of the three-dimensional z-stack to ensure that only nuclear microtubules were selected. 53BP1 foci were counted using the Spots MatLAB Xtension of Imaris or Difference of Gaussians (DoG) and Analyze Particles functions on ImageJ/FIJI. To determine the association of 53BP1 foci with LMNB1 tubules, the foci were three-dimensionally reconstructed using the Spots MatLAB Xtension in BitPlane Imaris 9.7 with the Different Spot Sizes option turned on to ensure capturing the actual size of each focus, followed by scoring the number of foci at or away from the tubules using the Find Spots Close to Surface MatLAB Xtension. Threshold was set to a value smaller than the average diameter of voxels at 0.3 μm, which was determined based on the average diameter of the 53BP1 foci (0.70 ± 0.21μm, total of 1117 randomly selected foci) and the average diameter of 110 randomly selected U2OS cells (13.0 ± 2.8 μm).

For nuclear zone-specific protein colocalization analysis, cells were divided into concentric zones of equal volume by creating nuclear shells of equal distance from the nuclear center normalized to the smallest nuclear volume value using BitPlane Imaris 10.0. Briefly, nuclei were selected by creating a new surface based on DAPI signal. The nuclear centers were determined by applying the Center of Mass to Spots Matlab Xtension. Afterwards, the Distance Transformation Matlab Xtension was calculated on the determined nuclear center result by selecting the default Transformation value of 1 to transform pixel sizes into distance measurements inside selected nuclei. Nuclei were then segmented into zones of equal distance from the nuclear center towards the periphery. The nuclear zones were masked by the DAPI channel to ensure that further analysis is performed exclusively inside the selected nuclear zone. To determine colocalization in different zones, the BitPlane Imaris Coloc function was then applied onto selected channels by using a threshold value of 0.2, and masking the overlapped channels with areas of created nuclear zones. The Summary statistics values representing intensity values of each protein of interest for single nuclei per nuclear zone were created. Those single nuclei protein intensity values were normalized over the smallest volume value of the nuclei’s respective nuclear zone.

For the FokI-DSB reporter system, image analyses were as follows. Colocalization of the single-DSB foci with 53BP1 foci in 2-6-5 cells was performed in BitPlane Imaris 10.0 using the Spots function by setting the lowest spot diameter value to 2, followed by visual confirmation of colocalization in single cells and colocalization value data extraction. DSB mCherry foci shape and counts were determined using the three-dimensional single-cell analysis in BitPlane Imaris 10.0. The distance between the single DSB foci and the nuclear edge (marked via LMNB1 IF) was measured in ImageJ/FIJI^89^ using the Straight Selection Tool by assessing the shortest distance from the DSB to the nuclear edge in three-dimensions. Single-cell analysis of the colocalization between the single DSB foci and LMNB1 was determined from the superimposed mCherry-DSB and LMNB1 IF channels in ImageJ/FIJI. Nuclear envelope tubule width measurements were performed as previously described^90^. Briefly, 3D-confocal images of selected cell lines stained for LMNB1 were analyzed in ImageJ/FIJI by detecting nuclear tubes in the XZ plane and measuring tubule diameters using the Straight Selection Tool.

### CRISPR/dCas9-based visualization of DSBs

The ends of a single DSB were visualized by inducing the lesion in the reporter 2-6-5 cells. DSB induction in reporter cells via a 1h treatment with 1μM Shield-1 ligand (cat# AOB1848, AOBIOUS) and 1 μM 4-Hydroxytamoxifen (cat# S7827, Selleckchem) results in the recruitment of the LacI repressor fused to mCherry and the nuclease domain of endonuclease FokI to the Lac operator (LacO) array located on the 5’ end of the single DSB reporter locus. The 3’ end of the single DSB reporter locus was also independently visualized via transient transfection of GFP-dCas9 and sgRNAs targeting the beta-Globin transgene located downstream of the LacO array. Cells were transfected with the sgRNAs and GFP-dCas9-expressing plasmid as previously described^94^. Cells were first transfected with sgRNAs and a GFP-dCas9 expressing plasmid for 48 h and subsequently treated for 1 h with 1 μM Shield-1 ligand (cat# AOB1848, AOBIOUS) and 1 μM 4-Hydroxytamoxifen (cat# S7827, Selleckchem). Cells were transfected with non-targeting control sgRNA (True Guide Syn. SgRNA Negative Control, Non-Targeting, Invitrogen A35526; Lot: 2102041) or with three custom targeting sgRNAs (Invitrogen, Supplementary Tables 1 and 2).

### Engineering SUN1 and control chimeric proteins

The human eGFP-Sun1 full length (eGFP-Sun1) gene was synthesized and cloned into the pcDNA3.1+ plasmid (Invitrogen) using NheI and XhoI restriction sites. Full-length human Ku70 was fused to eGFP and human Sun1 that lacks the lamina domain (Sun1ΔN). This 4,905-nucleotide gene (Ku70-eGFP-Sun1ΔN) was synthesized and cloned into the pcDNA3.1+ plasmid (Invitrogen) using NheI and XhoI restriction sites. A SV40 NLS, a nuclear localization sequence, and SV40 is simian virus 40, was engineered between Ku70 and eGFP. To ensure protein flexibility, a (GGGS)_4_ linker was inserted between Ku70 and SV40 NLS. GGS linkers were also inserted between the SV40NLS and eGFP, and between eGFP and Sun1ΔN. Both SV40 NLS-eGFP-Sun1 and Ku70-eGFP-Sun1ΔN contain a silent mutation on amino acid 425 on Sun1 that confer resistance to a siRNA with target sequence: CCGTGTTGAACTGGGCAAGCA. Ku70-eGFP and eGFP-Sun1ΔN were generated by digesting the Ku70-eGFP-Sun1ΔN plasmid with HpaI & XhoI, and NheI & AflII, respectively. The oligos AACTGAC & TCGAGTCAGTT, and CTAGCGCCACCATGC & TTAAGCATGGTGGCG were annealed and cloned into the digested plasmid using T4 ligase for Ku70-eGFP and eGFP-Sun1ΔN, respectively.

### Coimmunoprecipitation

Cells were seeded 48 h prior to treatment with DMSO vehicle control or etoposide for 1h at 80% confluency. Cells were harvested by trypsinization and lysed for 1 h with RIPA buffer with protease inhibitor. Lysates were incubated for 30 min with Benzonase (cat# E1014-5KU, Sigma) and cleared by centrifugation for 10 min at 4°C. 1 mg lysate was pre-cleared with Dynabeads™ protein G (cat# 10004D, Thermo Fisher) for 1 h with rotation at 4°C. 50 µL of pre-cleared lysate was removed for input. 50 µL of Dynabeads™ were washed in IP buffer (10 mM Tris pH 7.4, 1 mM EDTA, 1 mM EGTA, 150 mM NaCl, 0.5% NP-40, 0.2 mM sodium orthovanadate, and protease inhibitor), and incubated with an antibody for LMNB1 (cat# 16048, Abcam), Sun1 (cat# 24568-I-AP, ProteinTech), or rabbit IgG (cat# ab171870, Abcam) for 1 h with rotation at 4°C. Antibody-conjugated beads were washed twice with IP buffer and incubated with the pre-cleared lysate for 2 h with rotation at 4°C. Beads were then washed five times with IP buffer and the antibody-protein complexes were eluted from the beads with 3X Laemmli buffer at 95°C for 5 min.

### Immunoblotting

Cells were lysed with RIPA buffer (10 mM Tris-HCl pH 8.0, 1 mM EDTA, 0.5 mM EGTA, 1 % Triton X-100, 0.1 % sodium deoxycholate, 0.1 % SDS 140 mM NaCl, protease inhibitor (Roche)) for 1h on ice and vortexing every 10 min. Lysates were cleared by centrifugation for 10 min at 4°C. Protein concentration was quantified using the Bradford assay (Bio-Rad) and samples were diluted to equal protein concentration with RIPA buffer. Samples were boiled in 3X Laemmli buffer (187.5 mM Tris pH 6.8, 6% w/v SDS, 30% glycerol, 150 mM DTT, 0.03% bromophenol blue), run on 8% SDS-polyacrylamide gels or 4-20% Tris-Glycine gradient gels (Invitrogen Novex), and transferred to nitrocellulose membranes. Membranes were blocked in 5% skim milk for 1h at RT and incubated overnight at 4°C with primary antibodies diluted in 1% skim milk. Membranes were washed three times with TBT+0.1% Tween-20 (TBST) and incubated with IgG secondary antibodies for 1h at RT. Membranes were then washed five times with TBST and incubated for 5 min with ECL substrate (Clarity Western ECL substrate, Bio-Rad) in the dark. Membranes were developed with the ChemiDoc™ imaging system (Bio-Rad).

### Metaphase chromosome spreads

MDA-MB-436 shCTL, shKIF50 or shKIFC3 doxycyline-induced knockdown cells were seeded in 6-cm plates. 48 h post-seeding, cells were treated with DMSO or 2 µM olaparib for 16 h. The cells were then treated with 0.1 µg/mL KaryoMAX™ Colcemid™ Solution (Gibco™, 15212012) for 3 h at 37°C and 5% CO_2_. Harvested cells were resuspended in pre-warmed 0.056 M KCl hypotonic solution for 20 min at 37°C. The cells were then pelleted and resuspended by the dropwise addition of 5 mL ice-cold fixative (3:1 methanol to glacial acetic acid). The fixed cells were incubated on ice for 1 h before centrifugation and resuspension in fresh fixative. Cells were incubated overnight at 4°C in fixative and then resuspended in fresh fixative two more times. Cells were then resuspended in 200-500 uL of fresh fixative and dropped on glass slides over a 70°C water bath. Slides were dried vertically at 37°C for 30 min. DAPI was added, and the spreads were imaged at 100X with a Leica DM4000B fluorescent microscope.

### Growth curves and colony-forming assays

To assess long-term growth and determine growth curves in standard cultures, experiments were conducted as previously described with modifications^95^. Briefly, for growth curves, 1x10^4^ MDA-MB-231 or 5x10^4^ MDA-MB-436 cells with either shCTL, shKIF5B or shKIFC3 induced by doxycycline addition were seeded in 6-well plates. Every 2-3 days, the cells were counted using the Beckman Vi-CELL XR Cell Viability Analyzer for 11 days. For long-term colony-forming assays, shCTL, shKIF5B, and shKIFC3 doxycycline-induced knockdown MDA-MB-231 (5x10^2^) or MDA-MB-436 (5x10^3^) cells were seeded in six-well plates and cultured for 10-21 days. Colonies were stained with crystal violet and scored with ImageJ/FIJI. To assess growth in PARPi, 5x104 MDA-MB-436 cells with either shCTL, shKIF5B or shKIFC3 induced by doxycycline were seeded in 12-well plates. The next day, the indicated PARPi concentrations or vehicle were added. Media was replenished after 4 days and the cells were counted after 7 days of growth using the Beckman Vi-CELL XR Cell Viability Analyzer.

### Mouse xenotransplantation

All procedures were approved by and performed in compliance with the guidelines of the Princess Margaret Cancer Centre and the University of Toronto Animal Care Committee guidelines. Equal parts of MDA-MB-436 cells (1.5 × 10^6^) and Matrigel (Corning cat: 354248) matrix were injected into inguinal mammary fat pads of 12–14-week-old NOD/SCID/IL2Rgamma^null^ (NSG) mice (Jackson Laboratory, strain 005557). Tumor volumes ((length × width^2^)/2) were monitored regularly using caliper measurements for 22 days. Mice were euthanized by CO_2_ inhalation when the tumor size reached the ethical endpoint, in accordance with animal care facility rules.

### Analysis of human tumors

Representative whole slide images of breast cancer specimens were obtained as follows: the cBioPortal (http://www.cbioportal.org) was used to search the TCGA PanCancer Atlas for breast tumours containing mutations in BRCA1, BRCA2, or PIK3CA. A list of potential cases was generated, and then the whole slide image was identified on the TCGA website (https://www.cancer.gov/tcga). In each case, a representative high-power magnification image was obtained from a representative area of the tumour (formalin-fixed paraffin-embedded sections; stained with Haematoxylin & Eosin; approximate magnification x400). Three representative areas were subsequently randomly selected for nuclear groove quantification; in each of these areas a total of 10 nuclei were examined for quantification of membrane clefting. Care was taken to avoid processing and tissue artifact (e.g., chatter).

### Bioinformatics

The Cancer Genome Atlas (TCGA) RNA-seq data were obtained from the Genomic Data Commons Data Portal (https://portal.gdc.cancer.gov) and corresponded to fragments per kilobase of transcript per million mapped reads upper quartile (FPKM-UQ).

Pediatric tumor data from the Therapeutically Applicable Research to Generate Effective Treatments (TARGET) project (cancer types: Wilms tumor (WT), rhabdoid tumor (RT), and neuroblastoma (NBL)). The Molecular Taxonomy of Breast Cancer International Consortium (METABRIC) data were obtained preprocessed and normalized from the cBioPortal (http://www.cbioportal.org/public-portal/). The data for chronic lymphocytic leukemia (CLLE) and malignant lymphoma (MALY) were obtained preprocessed and normalized from the ICGC Data Portal (https://dcc.icgc.org/). The HR and NHEJ pathway gene signatures were obtained from a curation study^96^ and overlapping genes between signatures were excluded from the analysis (HR = 43 and NHEJ = 11 genes. The altEJ signature has been previously described^97^. The normalized gene signature scores were based on single sample Gene Set Expression Analysis (ssGSEA) computed using the Gene Set Variation Analysis (GSVA) software package^98^. The forest plots and heatmaps were computed using the foresplot and circlize^99^ and ComplexHeatmap^100^ packages in R software. CRISPR-Cas9 loss-of-function screening data across established cancer cell lines were obtained from the Cancer Dependency Map (DepMap; https://depmap.org/portal/home/#/)^63^. For STRING network analysis^101^, an initial exploratory interaction network composed of a selected list of 92 proteins including DDR signaling kinases, nuclear envelope proteins, kinesin motors, and tubulin variants revealed three connected clusters. Based on these findings, a secondary network highlighting the connections between the nuclear envelope, DDR, and kinesins was created using the settings of full network, confidence, all active interaction sources, minimum required interaction score of 0.400, and a K-means clustering of 5.

### Statistical analysis

Statistical analysis was performed in GraphPad Prism Software using Student’s t-test, Mann-Whitney test, or one-way or two-way ANOVA with Tukey’s, Dunnett’s or Sidak’s multiple comparisons test best suited for the specific experimental design and type of datasets analyzed. Unless otherwise indicated, replicate information is as follows. The data were generated using the indicated number of biological replicates. For blots, images are representative of data obtained from two independent biological replicates. For microscopy, images are representative of phenotypes observed in at least two independent biological replicates and quantifications were based on at least fifty cells per condition per replicate from each of three independent biological replicates unless otherwise indicated.

### Code availability

All scripts used to analyse data are available upon request.

