## Supplementary information, tables, and figures for "DNA double-strand break-capturing nuclear envelope tubules drive DNA repair"

### SUPPLEMENTAL INFORMATION

#### Supplementary tables

#### Supplementary movie legends

#### Supplementary figure legends

#### Supplementary figures

**Supplementary Table 1. sgRNAs used in this study.**

| Name | Sequence |
|---|---|
| sgRNA1 | AUGGACGAGCUGUACAAGUC |
| sgRNA2 | CAAGAUCCGCCACAACAUCG |
| sgRNA3 | GUCGCCCUCGAACUUCACCU |
| sgNT | AAAUGUGAGAUCAGAGUAAU |

**Supplementary Table 2. siRNA sequences used in this study.**

| Target | Sense | Antisense |
|---|---|---|
| SUN1 | GUGUUGAACUGGGCAAGCATT | UGCUUGCCCAGUUCAACACGG |
| SUN2 | CGUAUGGUGCUUGGUAUUUTT | AAAUACCAAGCACCAUACGtc |
| KIFC3 | CCAAUGCUGUGACUUUCGAtt | UCGAAAGUCACAGCAUUGGtg |
| KIF5B | CCAAUGCUGUGACUUUCGAtt | AUAACUCCAAUUGCGGUUct |
| NUP98 | GGAUUGUUUGGAACCAGUUTT | AAGUGGUUCCAAACAAUCCtc |
| NUP153 | CUUUAUUUUCUGGCCAGCGTG | CUUUAUUUUCUGGCCAGCGtg |
| RNF8 | GAGGAUUUGGUGUCACAUAtt | UAUGUGACACCAAAUCCUCgt |
| RAD50 | GGCCUUUAAGUGAAGGAAAtt | UUUCCUUCACUUAAAGGCCaa |
| NBS1 (NBN) | GGAAAAACUGUGCCAUUCUtt | AGAAUGGCACAGUUUUUCCtt |
| BRCA1 | Target sequence:<br>CAACAUGCCCACAGAUCAA<br>CCAAAGCGAGCAAGAGAAU<br>UGAUAAAGCUCCAGCAGGA<br>GAAGGAGCUUUCAUCAUUC | N/A |
| CTL | Silencer™ Select Negative Control No. 1 siRNA (Cat# 4390843, Ambion) /<br>Target sequence (for BRCA1 experiment):<br>UGGUUUACAUGUCGACUAA<br>UGGUUUACAUGUUGUGUGA<br>UGGUUUACAUGUUUUCUGA<br>UGGUUUACAUGUUUUCCUA | N/A |

#### Supplementary movie legends

**Supplementary Movie 1. Three-dimensionally reconstructed nucleus showing etoposide-induced LMNB1 tubules from around the nucleus associated with DSBs.** The cell was treated with etoposide for 1 h, and z-stacks were obtained using super-resolution imaging. Nuclear slicing shows the LMNB1 tubules in the XY plane with associated 53BP1 foci followed by three-dimensional nuclear reconstruction separating the boundary (transparent/grey) and tubular (green) LNMB1 signal. 53BP1 foci at or away from LMNB1 tubules were color-coded magenta or cyan, respectively.

**Supplementary Movie 2. Three-dimensionally reconstructed nucleus showing DSB-associated LMNB1 tubules re-emerge following nocodazole washout and etoposide treatment.** The three-dimensional LMNB1 nuclear signal and associated 53BP1 foci were re-constructed in three dimensions. Nuclear slicing shows the LMNB1 tubules in XY, XZ and YZ planes with associated 53BP1 signal. Subsequent reconstruction of LMNB1 and 53BP1 surfaces shows DSB foci at or away from the LMNB1 tubules. The reconstructed boundary (transparent/grey) and tubular (green) LMNB1 structures are shown.

**Supplementary Movie 3. The natural reversal of etoposide-induced LMNB1 nuclear tubules in a live cell highlights the dynamic nature of these nuclear envelope structures.** The live cell was treated with etoposide for 1 h and shown is one XY z-plane representing slice 36 of 85 z-slices, which is deep into the nucleus. The LMNB1 tubule reversal process lasted approximately 15-24 min. See Supplementary Movie 4 for the chromatin time-lapse imaging of the same nucleus.

**Supplementary Movie 4. Chromatin re-organization during the natural reversal of LMNB1 tubules.** The live cell was treated with etoposide for 1 h and shown is one XY z-plane representing slice 36 of 85 z-slices, which is deep into the nucleus. Chromatin resealing in the wake of the reversing LMNB1 tubule (see Supplementary Movie 3 for matching LMNB1 signal) lasted approximately 15-24 min. Together, Supplementary Movies 3 and 4 highlight the coordinated dynamics of LMNB1 tubule reversal and chromatin reorganization inside the nucleus.

#### Supplementary figure legends

**Supplementary Fig. 1. Experimental approaches and additional characterization of etoposide-induced LMNB1 tubules.** (a) LMNB1 imaging, reconstruction, and measurement of tubule formation by calculating the ratio of the total LMNB1 surface area (boundary + tubular) divided by the total surface area of DAPI-marked DNA to reveal the LMNB1 tubules score or invagination ratio. Also, see Supplementary Movies 1 and 2 for visualization of the reconstruction approach and reconstructed nuclei. (b) Representative XY and XZ planes from three-dimensionally imaged live-cell nuclei revealing the LMNB1 tubules assessed to reveal the percentage of cells with tubules and their width. (c,d) Effect of nocodazole treatment on LMNB1 tubules as assessed by the percentage of tubule-positive cells (c) or tubular width (d). (e) Cell cycle stage profiling of cells subjected to the indicated treatments. No changes were detected within the short timeframe of our experiments. (f) Representative images and quantification from live cells showing the induction of SIR-tubulin-stained microtubules displacing chromatin following etoposide treatment, their reversal upon etoposide removal, and their prevention by nocodazole treatment. (g) Fixed cell immunofluorescence showing endogenous Tubulin-positive filaments that can infiltrate chromatin and are repressed by pre-treatment with nocodazole. (h) Effects of the microtubule polymerization inhibitors nocodazole (NOCO) and vinblastine (VINBL) on etoposide-induced LMNB1 tubules. Data from (c) are reshown in (h) as they were part of the same experiment and to facilitate communication. (i) Representative two-dimensional images of 53BP1 foci association with etoposide-induced LMNB1 nuclear tubules. (j) Co-immunoprecipitation showed an etoposide-induced interaction between endogenous 53BP1 and LMNB1, as well as a partial decrease in total LMNB1 levels. (k) Live-cell time-lapse imaging showing the natural reversal of an etoposide-induced LMNB1 tubule even in the presence of the DNA damaging agent. Note that the tubule almost fully reverses within 15 min. Also, see Supplementary Movies 3 and 4. (l) Treatment with the microtubule polymerization inhibitors nocodazole (NOC) and vinblastine (VINBL) induces 53BP1 foci further in the presence of etoposide. (m) Representative images of RAD51 foci associated with etoposide-induced LMNB1 tubules. Foci excluded from the tubules can also associate with the LMNB1 at the nuclear boundary. (n) STRING network analysis suggesting connections between nuclear envelope components (top), kinesins (bottom), and the major DDR kinases DNAPK (a.k.a. PRKDC), ATR, and ATM. (o-r) Effect of knocking down various DNA repair proteins indicated on baseline and etoposide-induced LMNB1 tubules. (a-m, o-r) U2OS cells were used. (c-f, h, l, o-p) Data are shown as the mean±SD; n=3 biologically independent replicates, two-way ANOVA with Tukey's (c,h,l,o) or Dunnett's (d-f, p) multiple comparisons tests.

**Supplementary Fig. 2. Differential subnuclear distribution of DSBs and the induction of LMNB1 tubules under different experimental conditions and in diverse cell types.** (a-b) Colocalization analysis of the DNA damage marker γH2AX with either the KU70 NHEJ factor (left) or the RAD51 HR factor (right) in each of three concentric nuclear zones (C-center, M-middle, P-periphery) of equal volume in cells treated with vehicle or etoposide (ETP). (b,c) Levels of γH2AX (b) and LMNB1 tubules (c) in cells treated with indicated etoposide concentrations. (d) Representative images of rare nuclear defects observed upon etoposide treatment. (e) Etoposide increases the relative levels of cells with LMNB1 tubules in U2OS, HeLa, and the non-cancerous IMR90 cells. (f) Etoposide induces LMNB1 tubules in U2OS, HEK293T, HeLa, IMR90, and MCF10A cells. (g-h) Total 53BP1 levels and LMNB1 tubules are induced by treatment with the

DNA damaging agent etoposide, the microtubule inhibitor nocodazole (NOC), RNA Polymerase II inhibitor flavopiridol (FVP) or proteasome inhibitor MG132. (a-h) U2OS cells unless otherwise indicated; data are shown as the mean±SD; n=3 (a,b,f,g), n=4 (c), and n=9 (h) biologically independent replicates, two-way ANOVA with Tukey's multiple comparisons test (a), one-way ANOVA with Dunnett's multiple comparisons test (b,c,g,h), and multiple Mann-Whitney tests (f).

**Supplementary Fig. 3. Immunoblot controls for knockdowns.** Shown are immunoblots indicating the knockdown of SUN1, SUN2, NUP153, NUP98, KIF5B, and KIFC3 using small-interfering RNAs (siRNA). Actin served as loading control.

**Supplementary Fig. 4. Additional controls related to the FokI-DSB system.** (a) Schematic illustrating the ER-mCherry-LacI-FokI-DD (FokI-DSB) reporter system. (b) Colocalization of the induced FokI-DSB with 53BP1. (c) FokI-DSB foci association with LMNB1 tubules and with LMNB1 at the nuclear periphery. These data are reshown in (Fig. 3k) as they were part of the same experiment and to facilitate comparison. (d) Confirmation of siRNA-mediated protein knockdowns using immunofluorescence. (e) Knockdown of SUN1 or SUN2 partly decreased the percentage of tubules-positive cells following FokI-DSB induction. (f) Cell cycle stage profiling of cells following the knockdown of indicated factors. No statistically significant changes were detected within the short timeframe of our experiments. (g) Treatment with the histone deacetylase inhibitor SAHA is not redundant with SUN1 knockdown in terms of increasing the distance between FokI-DSB and the nuclear edge. (h-i) Visualizing one end of the FokI-DSB (h) using sgRNAs (sgDSB) and GFP-dCas9 expression (i) confirmed that the long and split FokI-DSB shapes represent less connected DSB ends. A non-targeting sgRNA (sgNT) served as control. (j-l) Enoxacin enlarges the 53BP1 foci surrounding the FokI-DSB (j,k) and fully restores the colocalization of the FokI-DSB with 53BP1 (l). (a-l) 2-6-5 cells; data are shown as the mean±SD; n=3 (c,d,g,k,l), n=2 (e), n=4 (f), and n=9 (h) biologically independent replicates; unpaired t-tests (c), two-way ANOVA with Tukey's multiple comparisons test (f,g), Mann-Whitney test (k), or two-way ANOVA with Dunnett's multiple comparisons test (l).

**Supplementary Fig. 5. Chimeric SUN1 fusion protein constructs and changes to the expression of endogenous SUN1 and SUN2 following etoposide treatment.** (a) SUN1 protein schematic showing the N-terminal lamina/chromatin-binding domain (magenta) and the C-terminal SUN domain (grey). Also shown are the confirmed post-translational modification sites of SUN1 from the UniProtein (middle) and PhosphoSite (bottom) databases. (b) Immunoblot showing decreased endogenous SUN1 protein levels upon siRNA-mediated knockdown. (c) Immunoblot demonstrating expression of the various SUN1 and control chimeric fusion proteins. Proteins of expected sizes are below the red asterisks. (d) Schematic of the various fusion proteins. (e) Immunoblot showing decreased levels of all endogenous SUN1 isotypes following siRNA-mediated knockdown. The differential reduction should be noted in different bands. (f-g) Immunoblots of endogenous SUN1 (f) and SUN2 (g) proteins from vehicle or etoposide-treated cells. (h,i) Etoposide treatment increases the phosphoserine signal (h) but the ubiquitin signal (i) co-immunoprecipitating with SUN1 (SUN1 IP). The blots in (h) and (i) are part of the same experiment but are shown separately to minimize the length of this figure.

**Supplementary Fig. 6. KIF5B and KIFC3 coexpression with defined genes and DDR pathway signatures across cancers.** (a,b) Heatmap showing unsupervised hierarchical clustering of PCCs between *KIFC3* (depicted violet) or *KIF5B* (blue) expression and the indicated genes coding for NHEJ factors (*TP53BP1*, *XRCC5, XRCC4*) or *LMNB1* (a), or the indicated genes coding for proteins representing different DNA repair pathways (NHEJ: *TP53BP1*, *XRCC6*; HR: *RAD51*, *RBBP8*; altEJ: *POLQ*, *LIG3*) (b), across cancer types (acronyms; BRCA indicates breast cancer; tumor numbers are shown). Nominal significant PCCs are indicated by asterisks (inset). (c-e) Forest plots of *KIFC3* or *KIF5B* coexpression analysis against NHEJ (c), HR (d), and altEJ (e) pathway signatures. The expression of *KIFC3* and *KIF5B* negatively and positively correlated with the NHEJ pathway signature more than the HR or altEJ pathway signatures across cancers, respectively. Significant negative and positive PCCs are depicted in green and red data points, respectively. Grey data points are non-significant.

**Supplementary Fig. 7. Tumor sections from breast cancer patients show nuclear tubule-like structures.** (a,b) Human breast cancer cases exhibit nuclear grooves. (a) Representative images of cases of human breast carcinoma with indicated mutations retrieved from the TCGA (The Cancer Genome Atlas; https://portal.gdc.cancer.gov/; see Methods). White arrowheads highlight nuclei showing nuclear grooves reminiscent of the nuclear envelope tubules uncovered herein in cell culture. (b) Quantification reveals the number of grooves per nucleus in different human breast cancer cases (n=10 cases for each of the *BRCA1* and *BRCA2* mutants). (c-e) Immunoblots confirm the knockdown of indicated factors in MDA-MB-231 or MDA-MB-436 cells. (f) Effects of KIF5B or KIFC3 knockdown on the formation of radial-like chromosome structures in MDA-MB-436 cells treated with vehicle or PARPi (2 μM, 16 h). The percentage of radial-like structures is shown (n=50 metaphase spreads). (g) Synthetic lethality effect or score for the indicated genes when combined with BRCA1 wild-type (n=68) or mutant (n=8) breast and ovarian cancer cell lines. CRISPR screen data accessed through DepMap (see methods) are presented as mean±SD; Mann Whitney test.

**Supplementary Fig. 8. Model for dsbNETs formation, structure, and function.** (**a**) Upon the induction of endogenous or exogenous DNA damage, the DDR signaling kinases mediate the formation of dsbNETs. The dsbNETs infiltrate the nucleus and are dependent on microtubules, LINC proteins SUN1 and SUN2, NPC basket component NUP153, and microtubule plus-end-directed kinesin-1 KIF5B. The dsbNETs are transient, and their reversal is mediated by the microtubule minus-end-directed kinesin-14 KIFC3. The dsbNETs bring the nuclear envelope and its resident proteins to DSBs internally located within the nucleus, providing the solid support necessary to reconnect the ends of a DSB to each other. (**b**) Under standard conditions, dsbNETs promote faithful DSB repair, avoiding chromosome (Chr.) defects and ensuring cell survival. (**c**) In BRCA1-deficient breast cancer cells treated with PARPi, the capacity of these life-saving drugs to kill cancer cells is mediated in part by dsbNETs.

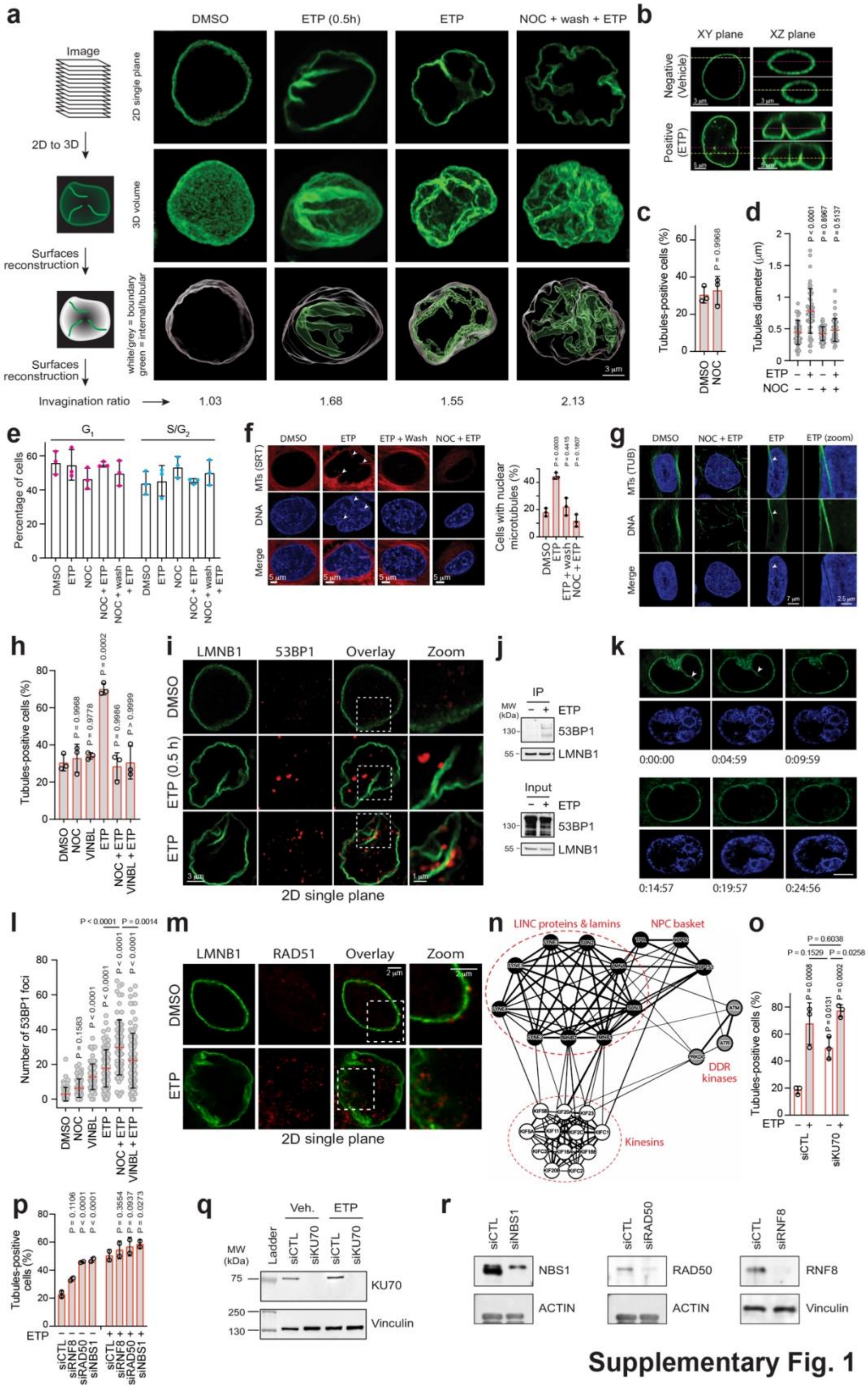


**Supplementary Fig. 1**

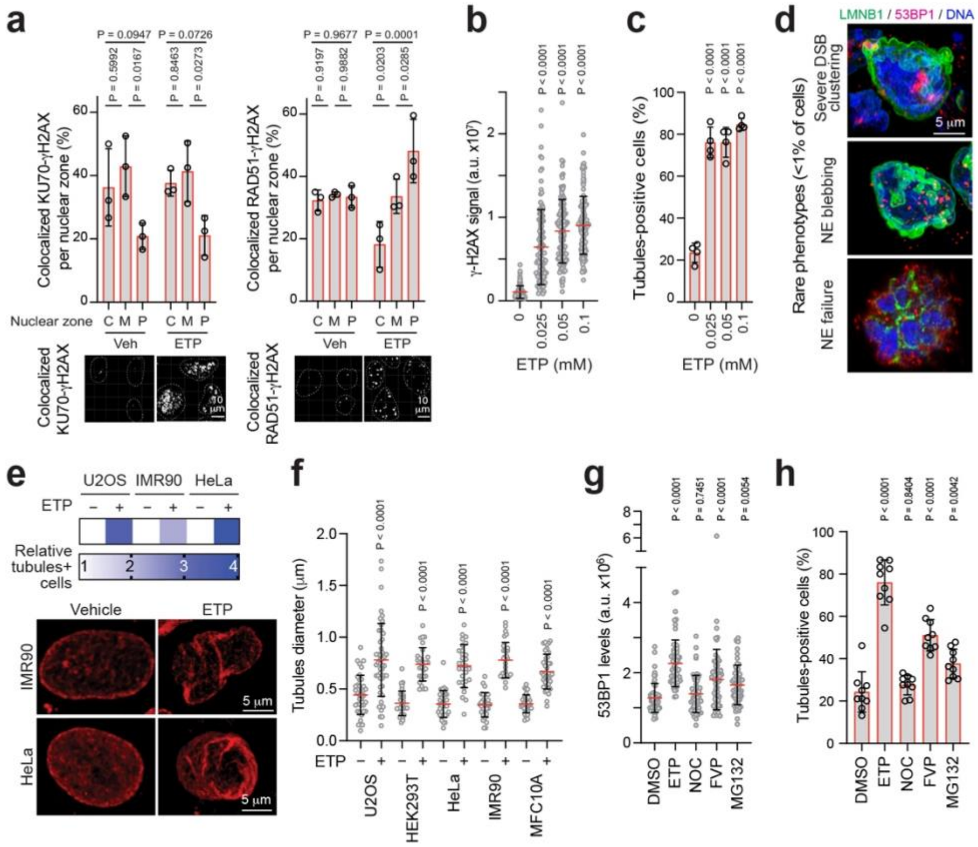


Supplementary Fig. 2

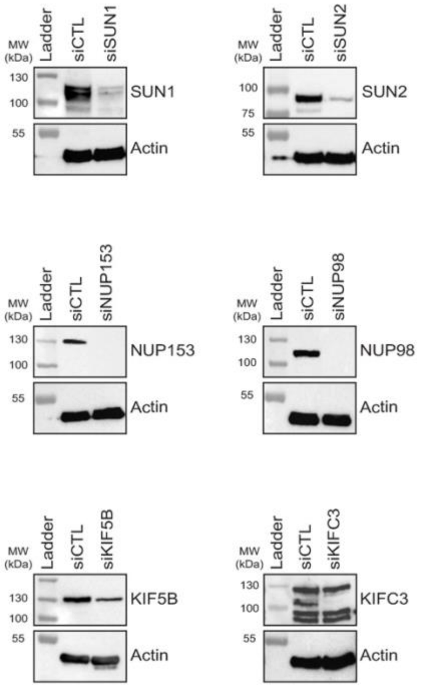


**Supplementary Fig. 3**

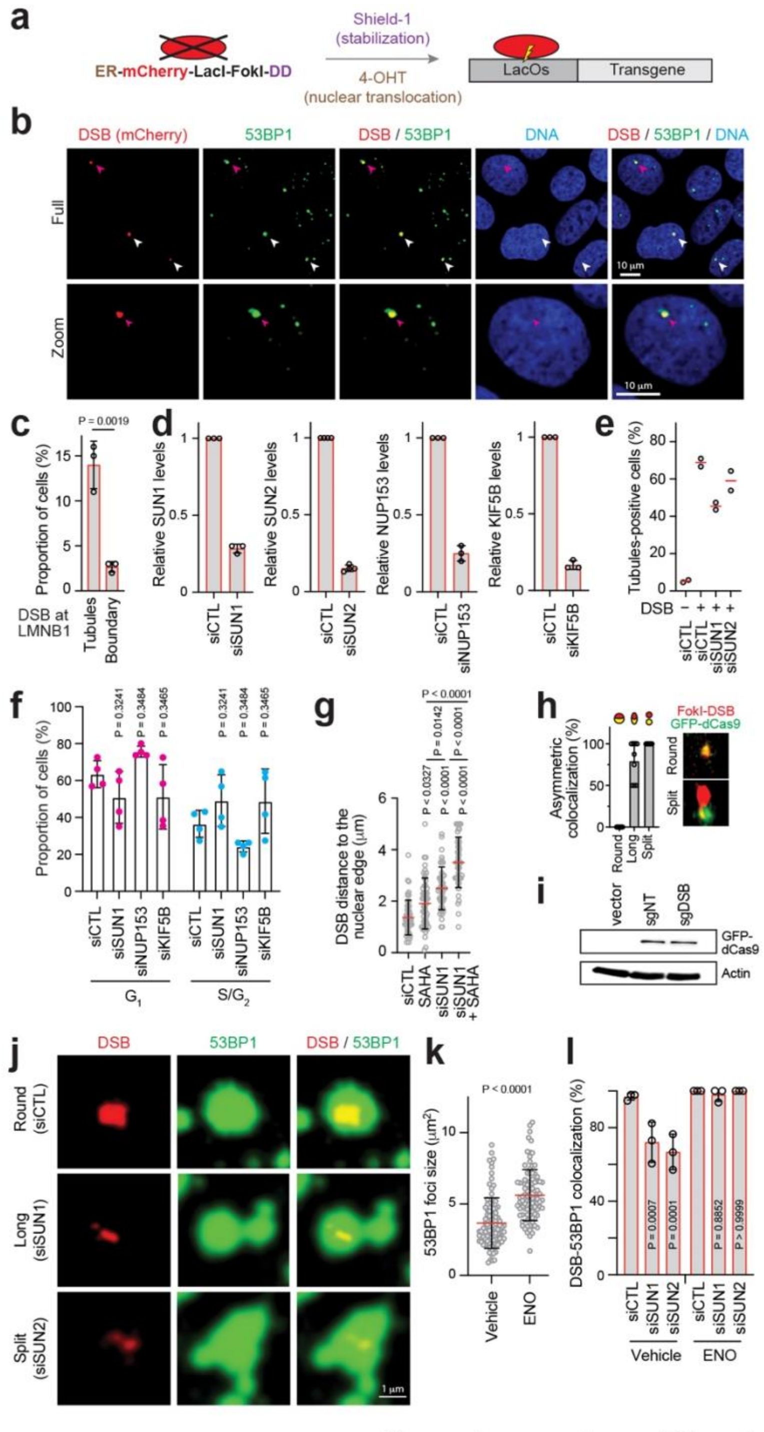


**Supplementary Fig. 4**

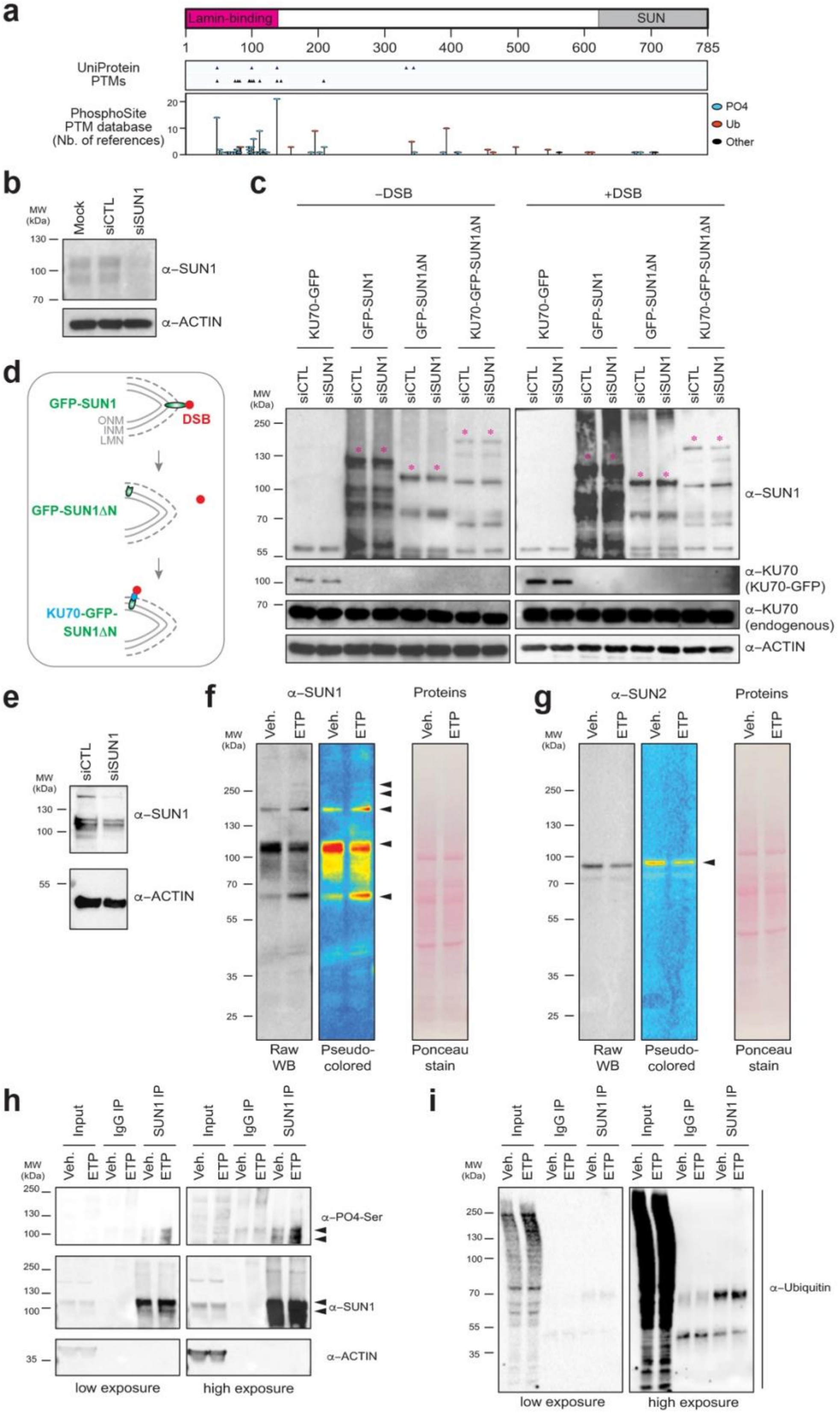


Supplementary Fig. 5

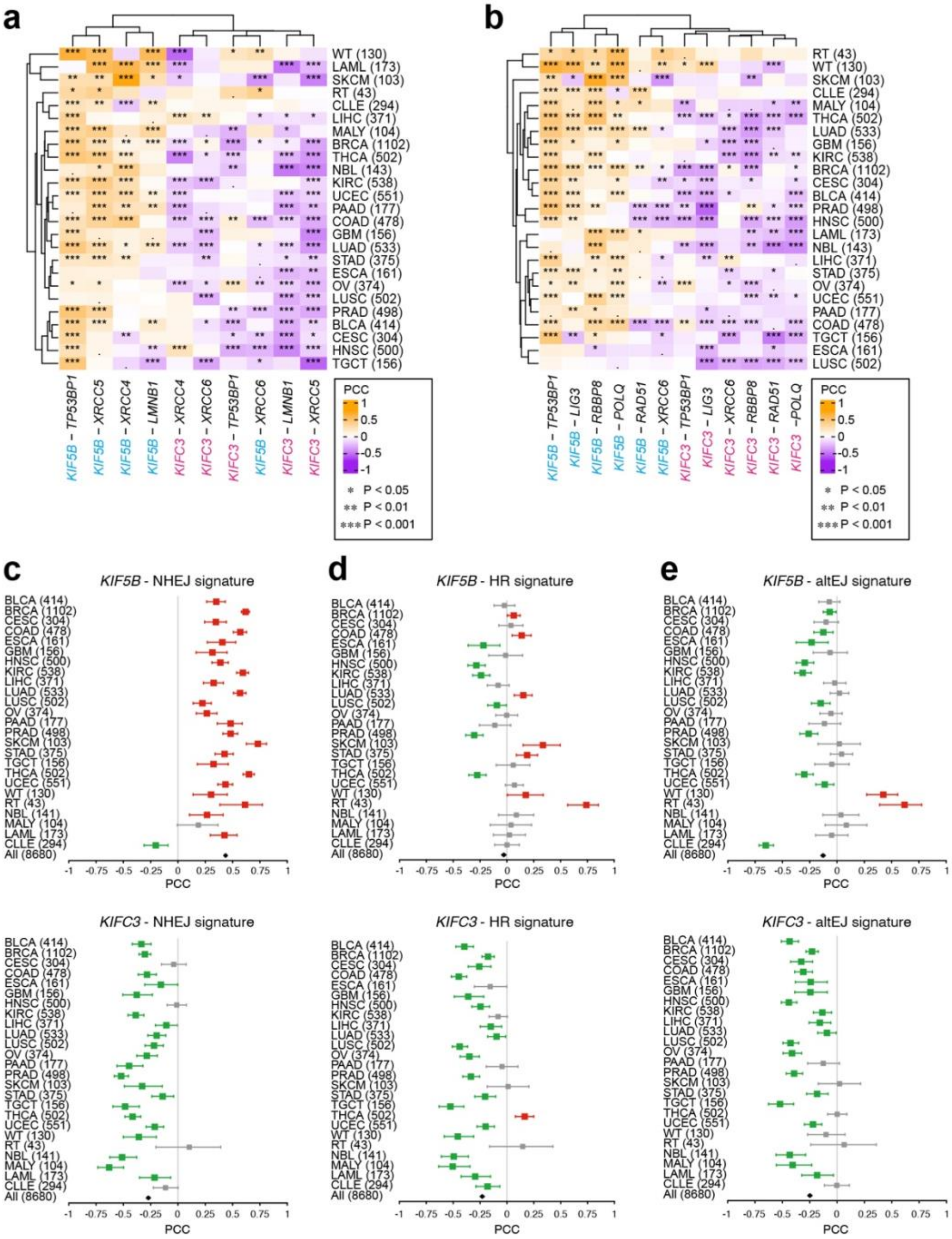


Supplementary Fig. 6

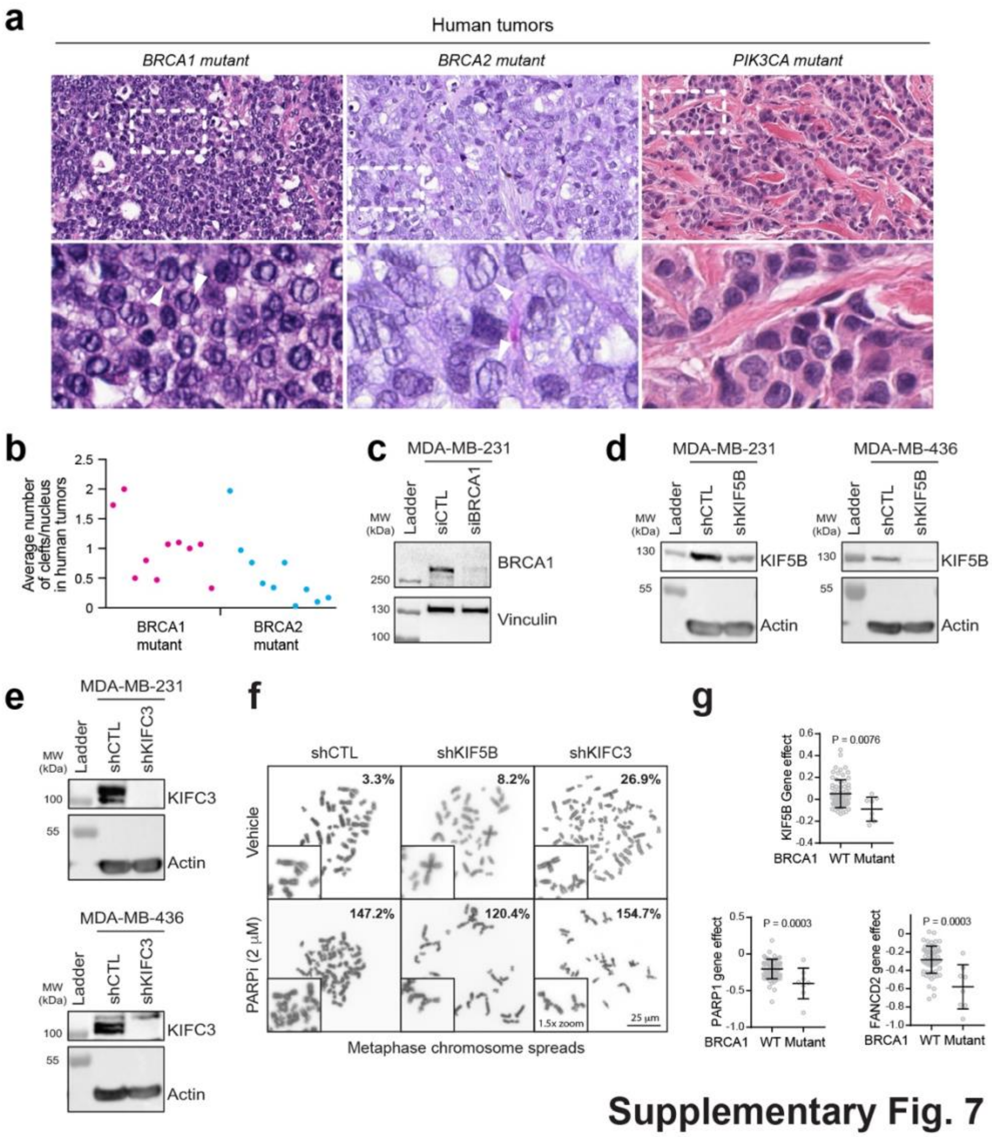


**Supplementary Fig. 7**

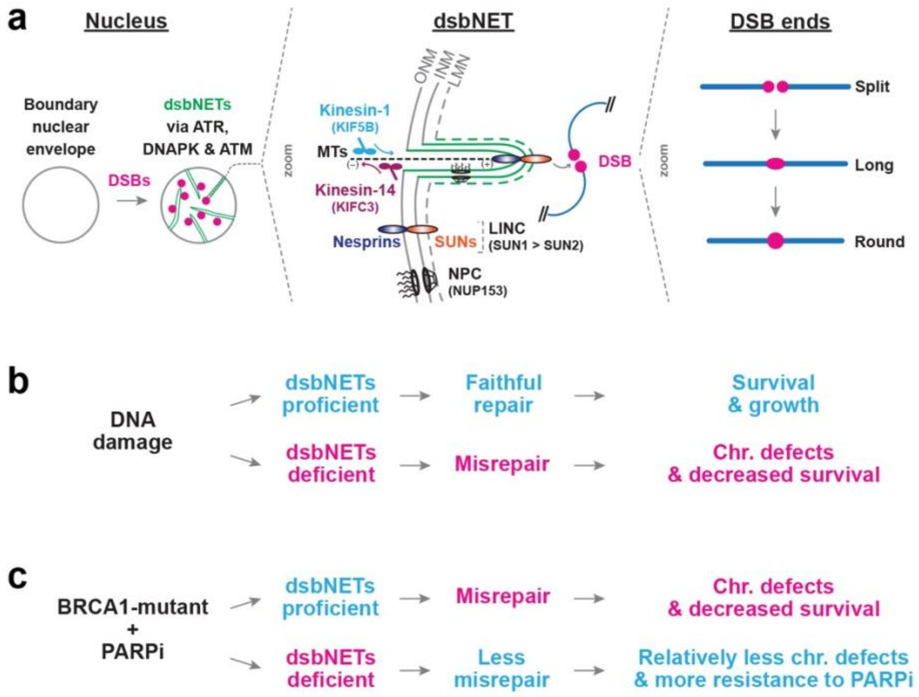


**Supplementary Fig. 8**
